# Estimating distances to desertification points from dryland ecosystem images

**DOI:** 10.1101/2024.02.20.581244

**Authors:** Benoît Pichon, Sophie Donnet, Isabelle Gounand, Sonia Kéfi

## Abstract

Resource-limited ecosystems, such as drylands, can exhibit self-organized spatial patterns. Theory suggests that these patterns can reflect increasing degradation levels as ecosystems approach possible tipping points to degradation. However, we still lack ways of estimating a distance to degradation points that is comparable across sites. Here, we present an approach to do just that from images of ecosystem landscapes’. After validating the approach on simulated landscapes, we applied it to a global dryland dataset, estimated the distance of each of the sites to their degradation point and investigated the drivers of that distance. Crossing this distance with aridity projections makes it possible to pinpoint the most fragile sites among those studied. Our approach paves the way for a risk assessment method for spatially-organized ecosystems.

---

The dynamics of complex systems are knowingly difficult to predict. Examples from past societies (*1*), climatic (*2*), or financial (*3*) systems emphasize that some complex systems can undergo abrupt transitions from their current state to a contrasting, possibly degraded one, once they pass a tipping point. In ecology, there are evidence of such abrupt transitions in drylands, savannas, or coral-reefs for example, where drastic changes in species composition and ecosystem functioning can occur with important ecological and social consequences (*4–6*).

Because these abrupt transitions are often unexpected, previous research has been devoted to the search for indicators of their approach. Generic ‘early warning signals’, such as temporal autocorrelation and variance, have been developed in theoretical models (*7*) and quantified on time series data from a variety of systems, including financial or societal crisis (*3*), emotion fluctuations in humans (*8*) and changes in ecosystem states (e.g., (*9–11*)). These temporal early warning signals, however, require long time series and can fail in a number of contexts (*12–14*), including in strongly spatially structured ecosystems (*15*).

In addition to exhibiting abrupt transitions, a number of ecosystems, such as drylands, mussel-beds, and salt marshes, also have a characteristic spatial structure (*16–18*). In these resource-limited environments, facilitative interactions between species and their local environment generate positive feedback loops that enhance the local environmental conditions and buffer species against disturbances (*16–19*). Such facilitation mechanisms, which are known to lead to abrupt ecosystem transitions, generate an emergent spatial structure, referred to as ‘self-organized’, with patches of individuals aggregated in space and interspersed by open areas (*20–22*).

These spatial patterns can be reproduced by dynamical models (*20, 23, 24*). In these models, when conditions become harsher, predictable changes in spatial structure are observed until the ecosystem switches to a degraded state, such as a desertified state deprived of vegetation in drylands. Studies suggested that, in self-organized ecosystems, spatial structure could hence be used as an indicator of resilience, meaning that it could inform us about the distance to the degradation point (*11, 25, 26*).

While spatial indicators of resilience provide a promising way of diagnosing a loss of resilience in a given system as it is approaching its degradation point, there is in practice not a single spatial metric used in the literature to quantify the spatial structure but a collection of them ; this means that, when considered jointly, some of them may provide conflicting information and that it is not obvious how to combine them into a single indicator (*14, 26, 27*). In fact, there is currently no methodology establishing an explicit link between the spatial structure and the resilience of self-organized ecosystems. We therefore lack the means to use spatial structure to quantify an actual distance to degradation points that is comparable across sites characterized by, for example, different vegetation, environmental, or soil conditions. Operationalizing these indicators to identify the most fragile ecosystem sites requires overcoming these limitations and finding an approach of estimating a distance to degradation points in a way that is comparable across field sites that differ in many ways (*28*).

## Estimating distances to degradation points

Here, we propose to tackle these limitations using an inverse approach that directly quantifies the resilience of a given ecosystem site, by estimating a distance to its degradation point from its observed spatial structure. The approach relies on a minimal, identifiable model which is able to generate the spatial structure typical of the self-organized ecosystem of interest, as well as the way the spatial structure changes along stress gradients (Methods).

Our minimal model relies on two parameters, namely *p* that quantifies the net recruitment of occupied sites (*e*.*g*. by vegetation) and *q* a spatial aggregation parameter related to the facilitation strength, which generates the emergent spatial structure. In order to infer these parameters from observations, we enhance this minimal model with an observational layer used to match the scales of the processes in the ecological model and in the observations, thereby allowing to work on ecosystem images independently of their spatial resolution (Methods).

We estimate the parameters of the minimal model from observed ecosystem landscapes using Approximate Bayesian Computing (ABC): we generate a large number of simulated landscapes from the minimal model, compare each observed landscape to simulated ones through a discrepancy measure based on the spatial structure, and retrieve a sample of parameters (*p*_*est*_, *q*_*est*_) that generate simulated landscapes whose spatial structures are the most similar to each observed landscape (Fig. 1A).

**Fig. 1:**
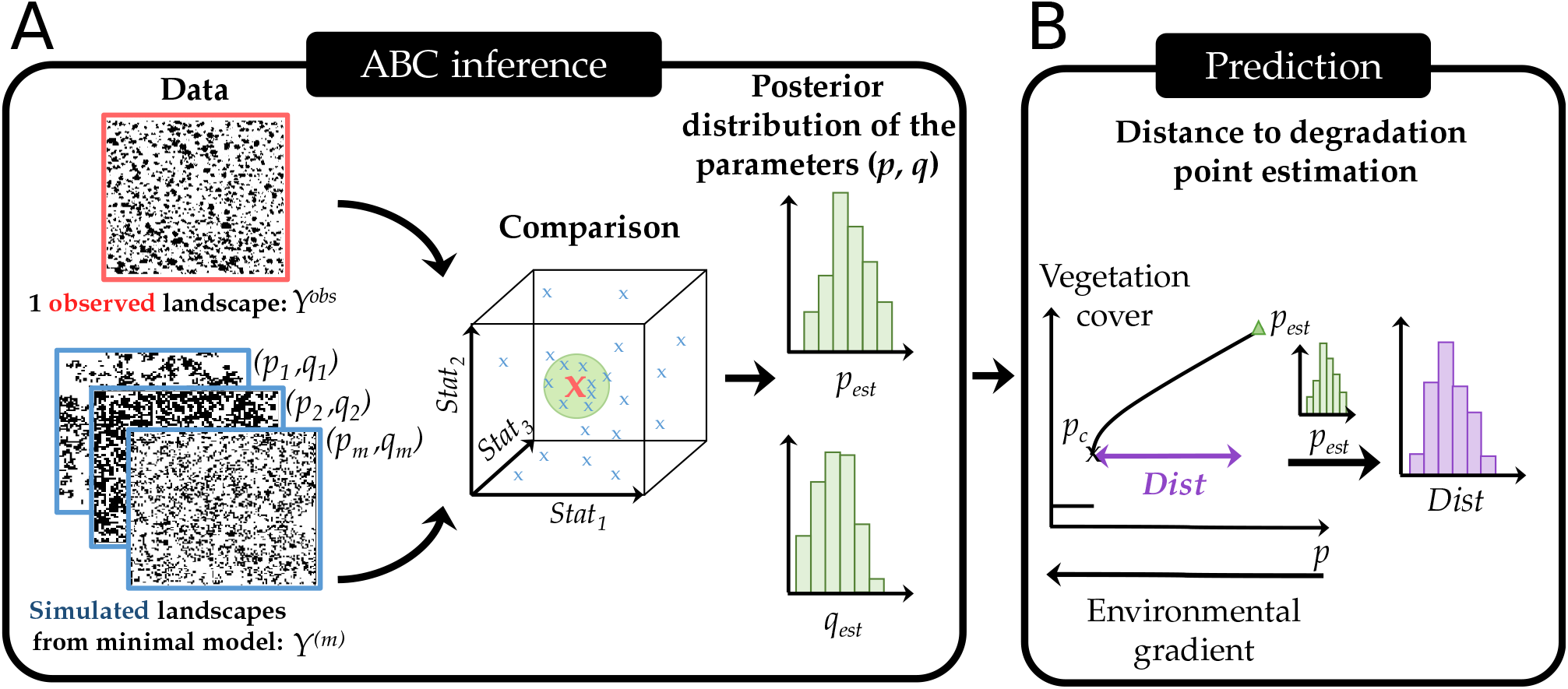
Overview of the inference approach to estimate distance to degradation point from spatially structured ecosystem images. The approach uses a simple mechanistic model (the so-called minimal model) and Approximate Bayesian Computing inference method to estimate parameters generating realistic spatial structures and derive distance to the degradation point for spatially structured ecosystems. (A) We linked observed (bottom row) and simulated (top row) vegetation landscapes using an inference approach (Methods) that allows estimating a distribution of the parameters of the minimal model ((*p*_est_, *q*_est_)). (B) Using the estimated parameters (*p*_est_, *q*_est_), we simulated a scenario of increasing abiotic stress and derived the distance to the desertification point in the minimal model.

For each observed landscape, once we know its estimated *p*_*est*_ and *q*_*est*_, we mimic a scenario of increasing stress by simulating the minimal model with a decreasing probability of recruitment *p* until the degradation point *p*_*c*_ is reached, below which the ecosystem shifts – either abruptly or gradually – to a degraded state deprived of occupied sites (Fig. 1B). This prediction procedure allows recovering the distance to the degradation point along the *p*-axis (*Dist* = *p*_*est*_ − *p*_*c*_; Methods).

### Validating the approach on simulated data

To test the robustness of the minimal model, we evaluated it on simulated landscapes from two models with different mechanisms than those of the minimal model but exhibiting similar emergent spatial patterns (Fig. S1). We used a dryland vegetation model with a more complex description of plant life history traits (seven parameters capturing dispersal, facilitation, recruitment, competition) (*23*) and a mussel-bed model containing three parameters (*17*) (Fig. 2A; Methods). Simulations from these two models generated a bank of observed spatial landscapes, for which the actual distance to the degradation point is numerically assessed. We then performed the approach on each of these landscapes and estimated a distance to the degradation point using the minimal model (Fig. 2B). Last, we compared this estimated distance to the ‘real’ one calculated in the two original models used to generate the simulated landscapes (Fig. 2C).

**Fig. 2:**
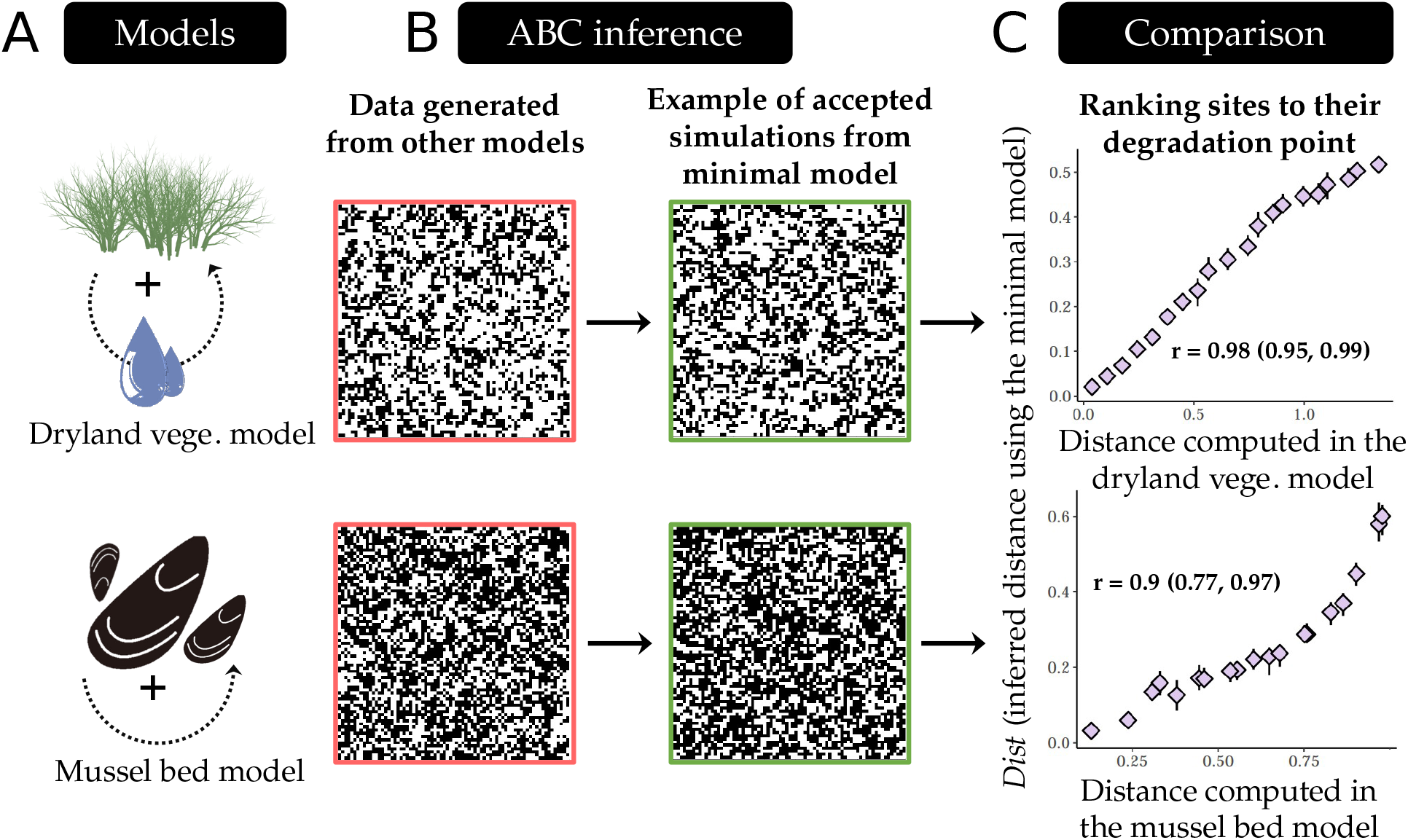
Validating the minimal model choice and inference approach using simulations. (A,B) We used a dryland vegetation model and a mussel-bed model that generate self-organized irregular patterns, and we performed the inference approach on landscapes generated by these models. (C) Then, we compared the distance to the extinction state computed in the two models (x-axis) to those estimated using our approach (y-axis). Spearman correlation is displayed on the panels with its confidence intervals. If our resilience estimation works well, the rank correlation between the distance to the degradation point predicted by our approach and the one computed in the two model is expected to be high (as observed here). In this case, it means that the minimal model is able to qualitatively ranking sites along a vulnerability axis defined as the distance to the degradation point (see Supplementary text for further discussion, and Figs. S2, S3 for a more extensive validation).

We find that the minimal model successfully reproduces spatial patterns generated using either the dryland vegetation or the mussel-bed model (Figs. 2B, Supplementary text). Importantly, the approach reliably ranks landscapes along an axis corresponding to the distance to the degradation point (median Spearman correlation = 0.98 and 0.9 for the dryland and mussel-bed models respectively; Figs. 2C and see S2, S3 for extensive parameter sets).

### Application to drylands

We next applied the approach to a global dryland data set corresponding to 293 images from 115 sites located around the globe. This allowed estimating the distance to the degradation point (here to the desertified state) for each of these images, making it possible to place each dryland image on a quantitative axis corresponding to a distance to the desertification point, and thereby providing an estimation of their resilience to upcoming perturbations (Fig. S4, and see as an example, the three sites in Fig. S5).

As expected, we find that sites with a higher vegetation cover tend to be farther away from their desertification point (Fig. S6). On the contrary, sites with a higher spatial autocorrelation of vegetation (Moran I) tend to be closer to their desertification point (Fig. S7A) in agreement with previous works (*4, 29, 30*). Importantly, quantifying the resilience of drylands allows to show that vegetation cover alone is not sufficient to predict the distance to the desertification point: the joint information of cover and spatial structure is needed to predict resilience (Fig. S8).

We used attributes measured at each site to investigate how local drivers affect directly and indirectly each of the three following response variables (Fig. 3): the resilience (distance to the desertification point), the vegetation cover, and the spatial autocorrelation of vegetation (Moran I). This allowed us to both check the coherence of the resilience estimations and to extract knowledge from these estimations. The local drivers considered were: the aridity level (1-precipitation/potential evapotranspiration), and a soil multifunctionality index measured as the average of the Zscores of 16 variables related to carbon, nitrogen and phosphorus in the soil (see Methods; (*19, 31*)). Higher values of soil multifunctionality have been related to more functional ecosystems (*19, 31*). These local drivers and response variables were all plugged into a structural equation model to disentangle relationships between drivers (Methods).

**Fig. 3:**
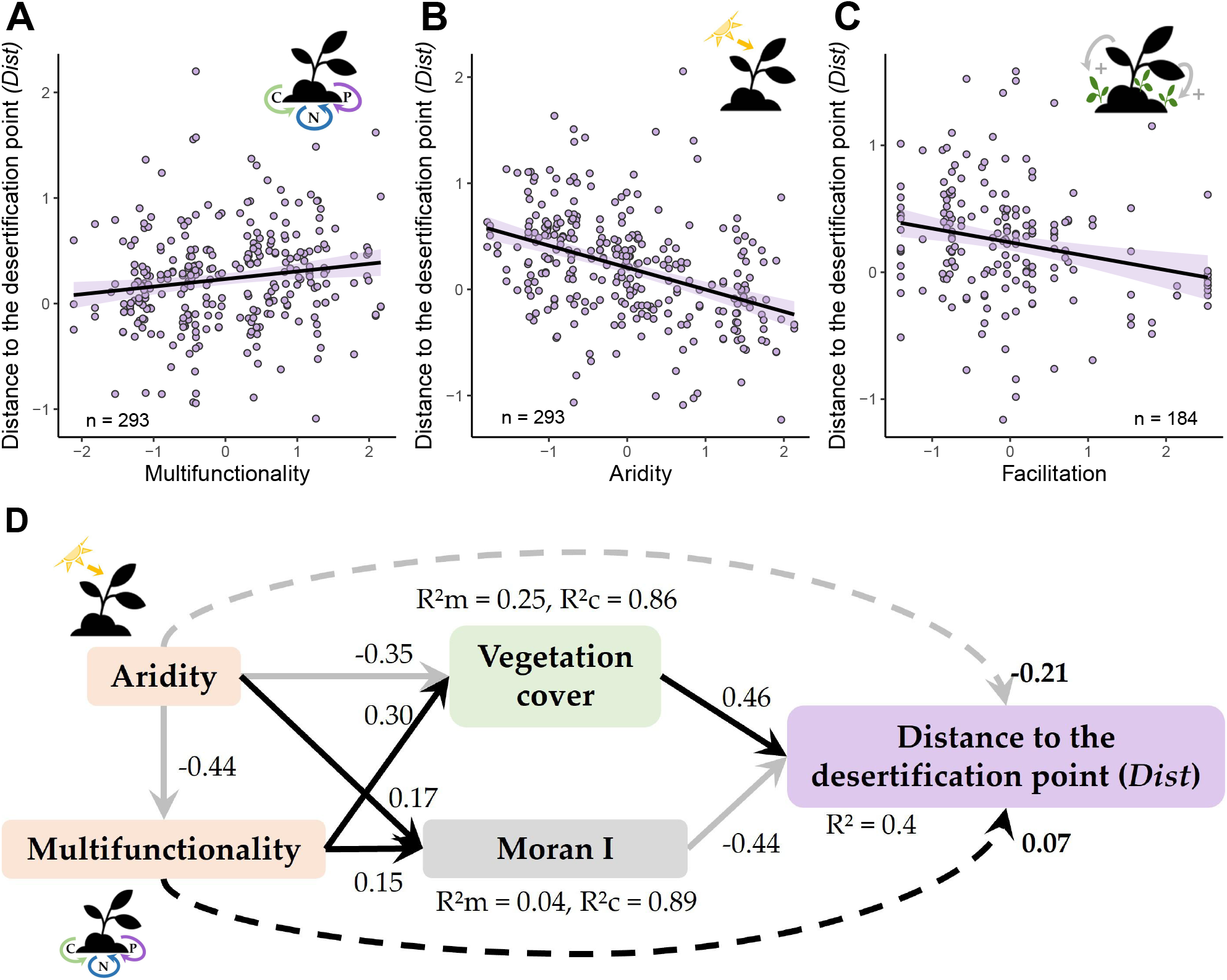
Estimating distances to the desertification points across global drylands. (A-C) Predicted responses of the distance of the desertification point (*Dist*) to changes in multifunctionality (A), facilitation strength (B), and aridity (C). The points show partial residuals, and the lines correspond to model fits. Shading around each line represents the 95% confidence interval. All slopes were significantly different from 0. Model fits are available in Table S1 and models *R*^2^ are available in Table S3. (D) Structural equation model decomposing the effects of aridity and multifunctionality on the vegetation cover, aggregation of the vegetation (Moran I) and the distance to the desertification point. Direct effects correspond to full lines, while indirect ones correspond to dashed ones. Facilitation was not included in the SEM since it was only measured for 70 sites among the 115 ones (184 landscapes). SEM: n = 293 landscapes, Goodness of fit: Fisher C stat = 1.584, degrees of freedom=2, *P-*value = 0.812. SEM coefficients confidence intervals are located in Table S2.

Consistently with previous studies (*16, 19, 22*), we found that sites with higher levels of soil multifunctionality are characterized by higher vegetation cover and spatial autocorrelation of their vegetation (Fig. 3D). This has been interpreted as reflecting the positive feedback between spatial structure, vegetation cover and ecosystem functioning (*19, 22*). Because vegetation cover and spatial autocorrelation have opposite effects on the distance to the desertification point (Fig. 3D), soil multifunctionality is found to be weakly positively associated with the distance to the desertification point (positive indirect effect in Fig. 3D), meaning that more resilient sites tend to show slightly higher functioning (Fig. 3A). By providing a comparable distance to the desertification point across ecosystem sites, these results complement previous knowledge on drylands by showing that the positive feedback between vegetation cover and ecosystem functioning may contribute to foster drylands’ resilience.

We replicated the same analysis with facilitation as an additional driver for a subset of sites for which the level of plant facilitation was measured in the field (i.e. *n* = 70 sites; Methods). We observe a negative association between facilitation and soil multi-functionality (Fig. S9) and a weak negative association between the level of facilitation and the distance to the desertification point, meaning that less resilient sites tend to show higher levels of facilitation despite a lower soil multifunctionality (Fig. 3C). This echoes the stress-gradient hypothesis, which suggests that the relative importance of facilitation compared to other processes such as competition increases with higher abiotic stress (*32*). Theoretical models suggest that higher levels of facilitation in plant communities buffer the effects of abiotic pressure and can lead to the maintenance of high vegetation cover for higher stress levels than without facilitation (*23, 33, 34*). At the same time, facilitation has also been shown to lead to possible abrupt transitions to desertification because facilitation contributes to positive feedback loops (*35*). So, as the stress level increases and communities approach their desertification point, increasing facilitation might maintain a relatively high cover along the gradient, but eventually lead to more abrupt collapses (from a higher vegetation cover) when communities reach their desertification point.

Last, because sites experiencing higher aridity levels have lower vegetation cover (Fig. 3D), this results in an expected negative association between aridity and distance to the desertification point (Fig. 3C,D). The negative effect of aridity on the resilience of the dryland sites is further mediated by the functioning of sites, since high aridity levels are associated with lower nutrient and carbon cycling (Fig. 3D). Given that drylands are experiencing increasing aridity levels (*36*), these changes will likely push drylands toward their desertification point, as it is already observed in some areas (*37*).

### Identifying the most fragile sites

We believe that our approach can provide operational information to pinpoint the most fragile sites of spatially structured ecosystems based on images. We illustrate a possible application of our approach by crossing our static resilience estimations for the dryland sites with aridity projections using the IPCC’s (Intergovernmental Panel on Climate Change) scenario assuming a sustained increase in carbon emission (RCP 8.5). We categorized the 293 landscapes into five vulnerability groups based on cluster analysis (Methods). Among these groups, the ‘high risk’ landscapes (red points in Fig. 4) are both relatively close to their desertification point and expected to undergo a rapid increase in aridity in the next decades. Some landscapes are projected to experience similar changes in aridity, but are further away from their desertification point, and therefore classified as under ‘climatic risk’ (orange and yellow points in Fig. 4). Compared to the high risk landscapes, these landscapes typically have a higher vegetation cover and slightly higher soil multifunctionality (Fig. S10). By contrast, landscapes under ‘ecological risk’ are relatively closer to their desertification point while projected to undergo relatively smaller increases in aridity in the future. Scaling-up such risk analyses to larger dryland areas and integrating future climatic and land-use changes would help targeting management toward the most fragile sites.

**Fig. 4:**
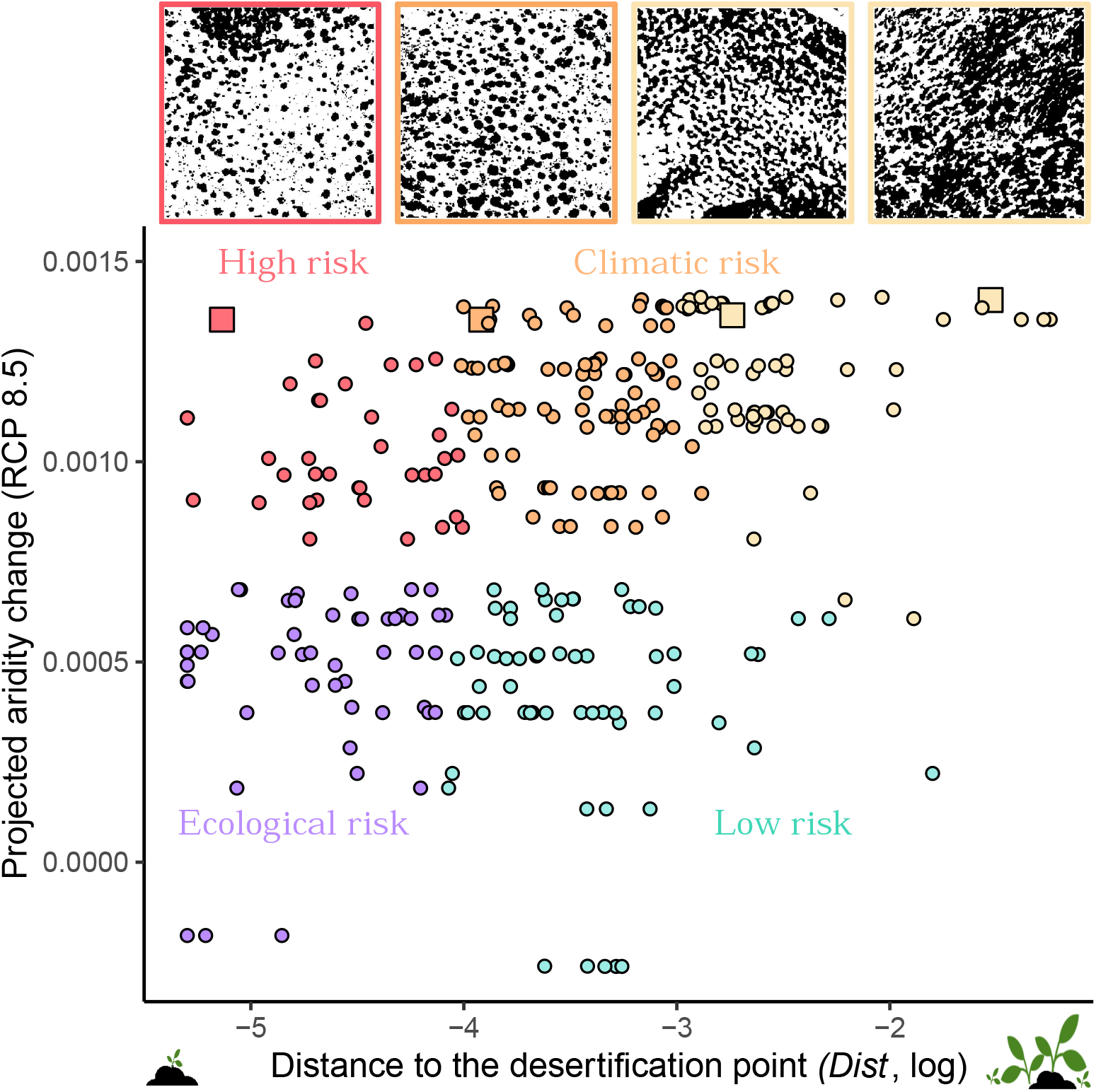
Crossing estimated distance to the desertification point with aridity projections to pinpoint most vulnerable sites. We mapped all sites into a 2D space defined by the distance to the desertification point (*Dist*, x-axis) and the linear trend of aridity for 1950–2100 under the IPCC RCP 8.5 scenario (y-axis). Then, we delimited four vulnerability clusters (“high risk” *n* = 38, “climatic risk” *n* = 51,”ecological risk” *n* = 144,”low risk” *n* = 60). Red points, for instance, correspond to sites both close to their desertification point and expected to undergo rapid aridity increase in the next decades. The characteristics of the clusters (current aridity level, multifunctionality, facilitation and vegetation cover) can be found in Fig. S10. Four vegetation landscapes located in Spain and expected to experience a rapid increase in aridity are given as examples. Their point shape is a square and is slightly bigger than the other points to allow their identification.

Drylands cover 41% of the land surface and host more than 2 billion people (*38*). There is thereby a need to build knowledge on these ecosystems and understand their possible responses to increasing levels of perturbations.

Our approach allows to recover the resilience (here estimated as a distance to a degraded state) of a given ecosystem site from a snapshot of its spatial structure. Previous studies have investigated how spatial indicators change along environmental gradients by comparing the spatial structure of different ecosystem sites in space (*11, 27, 39*); however, because the values of these indicators are not directly comparable with each other, this could not allow building resilience maps. Our approach provides a site specific estimate of resilience which allows ranking sites along a vulnerability axis defined as a distance to a degradation point. In other terms, spatial structure contains information about the response of a given site to future perturbations. This site-specific resilience complements studies proposing ecosystem risk assessment at larger scales using global thresholds (*e*.*g*., for aridity see (*40*)) and temporal information (*41, 42*), and therefore makes it possible to combine these different sources of information to build a more comprehensive assessment of ecosystem vulnerability (*43*).

Furthermore, our methodology is general enough to be applied to other spatially self-organized ecosystems. It relies on two key ingredients: *i*) identifying a minimal model informed from the mechanisms driving the ecosystem of interest which can reproduce the observed spatial patterns, and *ii*) coupling this minimal model to observed images from the ecosystem of interest. In such a case, because all images can be interpreted through the lens of a single minimal model, the model plays the role of a currency converter and allows estimating, for each site, a distance that is comparable across sites. The fact that the sites are now comparable makes it possible to map their resilience.

In fact, the minimal model we used can be relevant for other systems where entities aggregate in space because they are involved in local positive feedbacks that generate a characteristic irregular two-phase spatial structure (Supplementary text). This is the case, for example, in intertidal systems, where both mussels and sea-grasses perform facilitation, which promotes the emergence of self-organized spatial patterns (*17, 44*). When applied to a theoretical model of mussel-beds, our framework indeed proved to successfully predict the resilience of model intertidal ecosystems from their emergent spatial structure (Figs. 2, S3). It is noteworthy that the minimal model we used is nevertheless not suited for all ecosystems. In particular, we expect its applicability to be limited in cases where the ecosystem of interest is clearly characterized by multiple organism types, such as savannas where trees and grasses coexist. In such cases, the same inverse modelling approach together with a scale layer can be applied provided that a minimal identifiable model adapted to the system of interest can be determined. Incorporating the coexistence of different vegetation classes, would allow building a toolbox for estimating distances to degradation points in many spatially structured ecosystems.

It should be noted that our approach relies on estimating the distance to a degradation point using only one of the two parameters of our minimal model (the parameter *p*, which is the one that is assumed to best mimic aridity (see Methods). By boiling down the complexity of ecosystems to only two parameters, the resilience we estimate is in the unit of the parameter *p*, which allows putting all sites in the same reference system, thereby providing comparable estimations of resilience for all the sites. A drawback is however that the distance estimated cannot be easily translated into a timescale or a driver level (e.g. aridity or grazing). The use of long term data-sets (*45*) or historical images (*46, 47*) could help getting a better sense for the characteristic timescale of changes in drylands. Given that our approach represents an operational step towards comparing and ranking sites at risk in self-organized ecosystems, it may pave the way for reliable predictions of ecosystem resilience to environmental change.

## Materials and Methods

Our aim is to infer the parameters of a minimal model for an observed landscape with irregular patterns, and use the parameters of this fitted minimal model to estimate a distance of the observed landscape to a degradation point.

### Data: Observed vegetation landscapes

345 images of dryland ecosystems were taken from Berdugo et al., (2017) (*1, 2*). These images, which correspond to 50m × 50m landscapes, have a resolution sufficient to identify vegetation patches (spatial resolution lower than 0.3 meters per pixel). The 345 images are distributed over 115 geographical locations in 13 countries. The choice of theses geographic locations was guided by the BIOCOM dataset, which gathered different biotic and abiotic conditions on these particular locations (see (*1, 3*) for details; Fig. S11). For each geographic location, multiple soil attributes were collected: the soil sand content, the soil amelioration index (measured as the difference in organic carbon content between vegetated and bare soil areas) and the so-called multifunctionality index. This index is the average of the 16 soil variables associated with carbon, nitrogen and phosphorous cycling and storage, each one being individually centered and reduced (z-score) prior to averaging. A high value of multifunctionality means high values of many of the soil variables considered (but not necessarily all of them), and therefore a higher “functioning” of the dryland site in terms of recycling and storage of carbon and nutrients (see (*3*)). In addition, the level of facilitation (*i*.*e*., the proportion of plant species that are significantly more in the neighborhood of a facilitating species) was measured on a subset of 70 locations. To measure the latter, the number of individuals found in open areas is compared to those found under facilitating species (*i*.*e*., nurses) and an *χ*^2^ is computed on each pairwise interaction (see (*4, 5*) for further details).

#### Binarization of the images

Each image was transformed by Berdugo et al. 2017 to identify vegetated *versus* bare soil areas by performing a K-mean classification to partition the pixels in clusters of luminance intensity (using a monochromatic gray version of the image). The number of clusters was chosen to match the vegetation cover as measured in the field ((*1*) for details). From this K-means approach, a binary image of the presence and absence of vegetation was obtained. In the following, each binarized vegetation landscape of *N* × *N* pixels is denoted 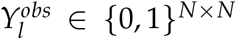, for *l* = 1, …, *L* (with *L* = 345 being the total number of landscapes).

### Minimal model of vegetation dynamics

We assume that each image of landscape 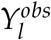 is the realization of a minimal model of vegetation dynamics at its asymptotic state. In this part, we drop the *l* index, since each landscape is treated separately. The minimal model used is hereafter referred to as the VDmin model (for vegetation dynamics minimal model).

#### Minimal model overview

The VDmin aims at mimicking stressed ecosystems such as drylands or salt marshes: in these ecosystems, some stress-tolerant plants facilitate their local environment by increasing water infiltration, providing shading, and increasing the local availability of nutrients (*6–10*). These mechanisms lead to the spatial aggregation of plants into patches separated by open bare soil areas (two-phase mosaic) (*11, 12*).

The VDmin model is a stochastic cellular automaton that describes the temporal evolution of a landscape composed of *C* × *C* (*C* = 100) cells that can be in two possible states: either colonized by vegetation (*V*) or empty (*E*). Each cell is assumed to correspond to the spatial scale of an individual plant (typically in drylands between 0.25*m*^2^ and 1*m*^2^). At each time-step, the states of the cells in the landscape change according to two probability parameters: *p* the local reproduction of vegetation, and *q* a spatial aggregation parameter that drives the spatial self-organization ((*13–15*); Fig. S12).

When *q* = 0, this model defaults to the Contact Process model (*13*) and only models growth and competition for space. By contrast, when *q≠* 0, the VDmin model also accounts for the spatial aggregation of plants because of facilitation. Preliminary analyses showed that, when *q* = 0, the model is not able to generate spatial patterns as the ones observed in the dryland data set investigated here. Therefore, as also suggested in other studies (*12, 14–16*), facilitation (as modelled by the parameter *q*) induces the emergence of irregular self-organized spatial patterns (although other processes can explain the emergence of spatial pattern, see (*17*) for instance). The importance of facilitation in the emergence of the spatial structure of drylands is a key assumption of our approach, which relies on the large empirical and theoretical literature emphasizing the importance of facilitation for the functioning and dynamics of drylands or other spatially structured ecosystems (*8, 12, 16, 18–22*).

Note that many other models can generate similar spatial structures (*e*.*g*., (***?***, *16, 23–27*)), because the minimal model used here belongs to a general class of models, where entities (whether it is vegetation, mussels, or people) tend to aggregate in space and form irregular patterns. Nevertheless, most of the other models are not identifiable, meaning that different combination of parameter values can lead to similar spatial structure. Our choice of this minimal model was therefore not only based on the ecological mechanisms but also on the identifiability of this model from static observations (see the “Parameter inference by Approximate Bayesian Computing” section), in addition to its ability to generate realistic spatial structures (see below).

#### Microscopic rules of the minimal model

The sequence of events corresponding to a time-step (from *t* to *t* + 1) in the minimal model is illustrated in Fig. S13. Formally, let 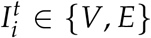 be the state of the cell *i* at time-step *t*, and let **v**(*i*) denote the four nearest neighbors of the cell *i*.

Starting from a landscape in state 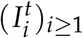 at time *t* and defining a random sequence of the vegetated cells in the landscape, iteration *t* + 1 corresponds to the following procedure applied on all the vegetated cells in the landscape, i.e.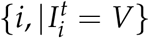.

1. Let *i* be the index of one vegetated cell in the landscape 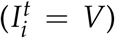, hereafter referred to as the focal cell (crossed cell in Fig. S13).
2. Let *i*^*′*^ be a randomly chosen cell among the neighbors of the focal cell *i* (*i*^*′*^ ∼*U*{**v**(*i*)}, where **v** are the neighbors of the cell *i* and *U* is the uniform distribution).
  - If the cell *i*^*′*^ is empty (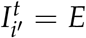, top row in Fig. S13), then either
    - it becomes colonized by vegetation with a probability 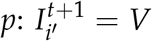
    - or the recruitment fails, and the focal cell dies with a probability 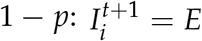
  - If the cell *i*^*′*^ is vegetated (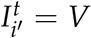, bottom row in Fig. S13), then
    - either a random cell in the neighborhood of the pair **v**(*i, i*^*′*^) becomes vegetated with a probability *q* (facilitation event,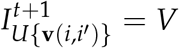)
    - or the focal cell dies with a probability 1 − *q* leading to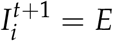.
  - Return to step 1 by considering the following vegetated state, until all cells have been taken one by one.

We ran simulations during a transient period of 1500 iterations, starting from a landscape of size *C* × *C* with high initial cover (randomly taking 80% of the cells initially colonized by vegetation). The 1500 time-steps transient period was enough to reach the asymptotic convergence of all simulations (preliminary analyses).

#### Phase diagram of the minimal model

Because the VDmin model includes a facilitation mechanism (through the parameter *q*), compared to the Contact Process model (when *q* = 0), the VDmin model exhibits both gradual and abrupt transitions to an empty (desert) state. To show this, for any parameter combination of *p, q*, we solved the mean-field approximation of the model, which assumes no spatial structure. The dynamics of the vegetation cover (*V*) using this mean-field approximation read as follows (Eq. S1):

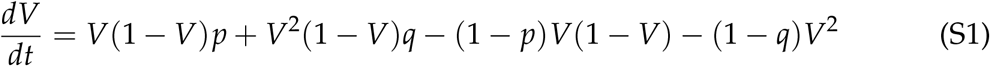

The first two terms correspond to recruitment or growth, while the two last terms correspond to plant mortality. By solving the minimal model for any *p* and *q*, we obtain a phase diagram that predicts both continuous (at low *q* values) and abrupt transitions (at higher *q* values) as illustrated in Fig. S14).

#### Modeling the scale of observation

As previously mentioned, the VDmin model is defined at the scale of the individual plant, and therefore a vegetated cell is assumed to correspond to one plant. However, the pixels of the observed images analyzed are generally at a finer spatial scale, thereby creating a mismatch of scales between the model and the observed vegetation landscapes (see Fig. S15 for illustration). To characterize the spatial structure of an image, we used spatial statistics (see “Choice of the discrepancy between observations and simulations” selection), which might be highly correlated with the spatial resolution of the image (*28–31*): for example, Fig. S16 shows that increasing the spatial resolution of the observed images keeps the cover and patch-size distribution unchanged but affects other statistics such as the spatial autocorrelation or the spatial variance.

To account for this mismatch of scales between the minimal model and the observed images, we added an observation layer *O* to the VDmin model. This layer involves one parameter *η* representing the scale of observation of a vegetation landscape generated from the minimal model. Practically, to account for this scale of observation, any cell of a realization of the model is replaced with a square of size *η* × *η* cells with the same type of pixel (vegetated or empty). Together with the scale layer, the VDmin has now 3 parameters (*θ* = (*p, q, η*)) which have to be estimated.

### Parameter inference by Approximate Bayesian Computing

#### The principles of ABC

The definition of the VDmin model combined with the scale layer makes the likelihood intractable. Approximate Bayesian Computing (ABC) has proved to be a convenient method for Bayesian parameter estimation when the model is easily simulated but has non-explicit likelihood (*32–35*).

The principle of the ABC, illustrated in Fig. 1, is to generate a set of parameters (*θ*^(*m*)^) = (*p*^(*m*)^, *q*^(*m*)^, *η*^(*m*)^) under the prior distribution (*θ* ∼ *π*). For each parameter *θ*^(*m*)^, a vegetation landscape *Y*^(*m*)^ is simulated under the VDmin model specified by the parameter *θ*^(*m*)^. If the simulated landscape *Y*^(*m*)^ is “too different” from the observed landscape *Y*^*obs*^, then the parameter *θ*^(*m*)^ is rejected, otherwise it is accepted. The simulated and observed landscapes are compared to each other through a discrepancy *D*(*Y*^(*m*)^, *Y*^*obs*^) defined in Equation (S2) and based on summary spatial statistics (see below). The ABC thus produces a set of accepted parameters (*θ*^(*m*)*∗*^) which will form a sample from an approximation of the posterior distribution uniform prior distribution:Note that we could have considered Monte Carlo Markov Chains methods, Sequential Monte Carlo, or parametric likelihood estimation (using an approximated version of the likelihood) that have also been proposed for parameter inference in spatial contexts (*34, 35*). However, the ABC method enables to decrease the computational cost of inference by using the same set of simulations for the inference of all images (*n* = 345; (*36*)).

#### Priors

The Bayesian inference relies on a prior distribution on the unknown parameters, namely (*p, q, η*). Because *p* and *q* are probabilities, we chose an empirical prior distribution ensuring that the resulting vegetation cover lies in [0.05, 0.9] (see the corresponding prior distribution of *p* and *q* in Fig. S17). The scale parameter *η* was sampled from a uniform prior distribution: *η* ∼ 𝒰_{1,…,5}_. The range for *η* was chosen following the result of a principal component analysis comparing observed vegetation landscapes with the simulated ones from the prior (Fig. S18). We sampled M = 367500 parameter.

#### Choice of the discrepancy between observations and simulations

A difficulty of the ABC relies on the choice of a similarity measure between the simulated and the observed vegetation landscapes. We built a discrepancy based on 11 spatial summary statistics summarizing the complexity of the spatial structure. The spatial statistics are the following ones:

- First, we computed the vegetation cover (*S*_1_) (fraction of vegetated sites), *S*_2_ the average number of plants in the neighborhood of each vegetation site (four nearest neighbors *i*.*e*. von Neumann neighborhood), as well as *S*_3_, the log-transformed vegetation clustering (average number of neighbors divided by the vegetation cover).
- In addition, we used four summary statistics that capture the level of vegetation aggregation: namely *S*_4_ the spatial variance, *S*_5_ the spatial skewness, *S*_6_ the near-neighbor correlation (Moran I), and finally *S*_7_ the log-transformed spectral density ratio (SDR; see (*37*) for details). *S*_7_ estimates the relative importance of long-range spatial variations compared to the total ones. We also computed *S*_4_ the spatial variance and *S*_5_ spatial skewness following (*38*). These metrics were computed from the R-package **spatialwarnings** (v3.0.3 ; (*39*)).
- Finally, we computed four metrics extracted from the patch-size distribution (PSD). In models of spatially structured ecosystems, patch size distributions have been shown to follow a pure power law at the percolation point (the point at which vegetation connects the landscape from one side to the other, (*25, 40, 41*)), and is known to start deviating from a power law to a truncated power law and then to an exponential law when the stress increases and cover decreases (*i*.*e*., as the system approaches its degradation point). Patch-size distribution has therefore been suggested to be a good spatial indicator of the resilience of dryland ecosystems (but also showed in mussel beds (*24*) or seagrass (*15*)). We therefore characterized the PSD by: *S*_8_, the power-law range (*i*.*e*., the fraction of the distribution of patch sizes fitting a power law (*1*)), and *S*_9_, the exponent of the power-law or truncated power-law which best fitted the patch size distribution. If an exponential law was the best fit for the PSD, we set the exponent value of the landscape to NA. In addition, we computed *S*_10_ the log-transformed fraction of the landscape covered by the largest patch (size biggest patch /total number of cells), as well as *S*_11_ the coefficient of variation of the patch size distribution. The latter allows accounting for the potential differences of heterogeneity in patch-size distribution between the model and the observed vegetation landscapes. We computed *S*_8_ to *S*_11_ using the functions *patchdistr sews* and *raw plrange*.

Note that, as illustrated in Fig. S19, these metrics are invariant with respect to the landscape size. Moreover, spatial statistics related to the patch-size distribution (*S*_8_ to *S*_11_) are invariant with respect to the spatial resolution of the vegetation landscape (Fig. S16). These summary statistics are logically correlated, as illustrated in Fig. S20, but we argue that they all bring important information to characterize the spatial structure of the landscape. When using ABC, we cannot ensure that different sets of spatial statistics would not increase the accuracy of our inference approach (*33*). Nevertheless, we made sure that spatial statistics were sufficient by (*i*) validating the identifiability of the model parameters (*p, q, η*) using simulated landscapes (below), and (*ii*) tested the robustness of the ABC method to different subsets of the spatial statistics used (see Fig. S21). We give in Fig. S22 the distribution of spatial statistics for simulations and observations.

To build a global discrepancy between observed and simulated landscapes taking into account the 11 spatial statistics, we first Box-Cox transformed and normalized them to be able to combine them, following (*36*). The obtained statistics are hereafter denoted *s*_*k*_.

Finally, the discrepancy between observed and simulated vegetation landscapes is the Euclidean distance between the respective summary statistics:

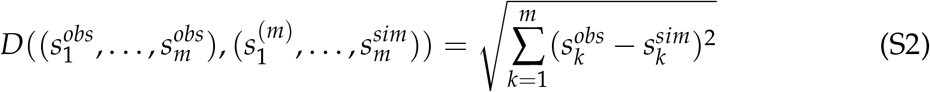

We followed the two-stage procedure proposed by (*36*) to select the 100 parameters, leading to the closest simulations. This is roughly similar to the fraction of simulations kept in (*42*) and a good trade-off between number of accepted simulations and the bias of the posterior parameter distributions (Fig. S23).

Note that we have considered post-sampling methods to reduce the bias of the posterior distribution (*32, 43, 44*). Post-sampling regression methods performed well on simulations, but none of them produced improvements on all the landscapes from the real dataset (as assessed by posterior predictive checks), so we kept the simple rejection algorithm without post-sampling regression. All the parameter inference by ABC was performed using the *abc* function from the **abc** R-package ((*45*), v2.2.1).

#### Illustration of the robustness of the inference method using simulated data

We investigated the robustness of our inference approach using simulated landscapes for which parameters are known (also called “virtual landscapes”). We tested the ability of ABC to retrieve the parameters of 100 virtual landscapes (each landscape corresponding to a randomly chosen pair of parameters (*p*_*l*_, *q*_*l*_)). For each landscape *l*, we simulated 5 scales of observation leading to *L* = 500 virtual landscapes, whose parameters were inferred by the previously described ABC procedure. We denote 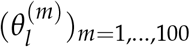 the posterior sample for landscape *l*. For each parameter *p, q* and *η*, we computed the so-called root mean-squared error (RMSE):

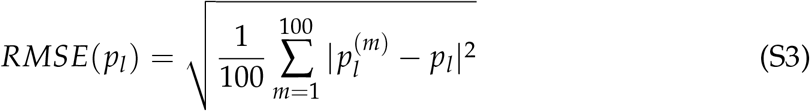

We normalized this metric by the RMSE computed under the prior distribution (NRMSE):

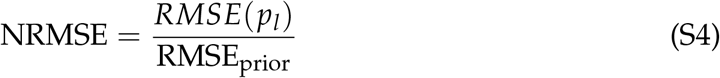

A NRMSE close to zero corresponds to a posterior distribution concentrated around the true parameters *p*_*l*_, while when it approaches one, the posterior distribution has not been updated using the data (*θ*^*∗*^ ∼ *π*). The same formula was used with *q* and *η* instead of *p*.

This analysis shows that we are able to infer parameters of simulated landscapes with high accuracy (mean NRMSE = 0.19 for *p*, mean NRMSE = 0.17 for *q* and mean NRMSE = 0.005 for *η*; Fig. S24). This high accuracy of inference for *p* and *q* is observed no matter the parameter *η*. In addition, we show that accounting for the scale of observation does not change the parameter that we infer for each virtual landscape (Fig. S25). Indeed, on average, we observe that the median of the posterior distribution of these 100 simulated vegetation landscapes is similar across all scales of observation (*η*). This analysis therefore shows that (*i*) the scale layer does not compromise the identifiability of the VDmin parameters, and that (*ii*) the parameter we infer for each site only depends on the spatial structure of the vegetation but not on the spatial resolution with which an image is characterized. All in all, this entails we can apply the ABC method to all observed vegetation landscapes **independently** of their spatial resolution.

#### Identifying a minimal set of spatial statistics

Since some spatial statistics are correlated with each other, it means that we could in practice use only a subset of all the spatial statistics while maintaining a good estimation of the parameters in the posteriors. In fact, the principal component analysis displayed in Fig. S12c shows that the first three principal components account for 90% of the total variance in spatial statistics, suggesting that the minimal required spatial statistics for the inference is smaller than 11. We therefore tested dimensional reduction of the number of spatial statistics used by performing a partial-least square analysis prior to the selection of the closest simulations using the AS.select function from the **abc.tools** R-package (*46, 47*). This projection technique consists in finding a set of orthogonal linear combinations of the original spatial statistics highly correlated to the parameters of interest (here the parameters *p* and *q*). In Fig. S21, we show that using a partial-least square analysis prior to ABC maintains a high accuracy of the posterior estimation, similar to what is obtained when using all spatial statistics (blue *versus* red points). We also illustrate the fact that considering a too small number of statistics by taking only the spatial statistics that are independent of the spatial resolution of the ecosystem image (see Fig. S16 for the ones kept) could lead to the degradation of the quality of the posteriors of the parameters (see Fig. S21, green *versus* red points). Since (*i*) computing the eleven spatial statistics is not computationally intensive because they are all derived from the same image, and (*ii*) that using all spatial statistics gives similar accuracy for the posterior estimation compared to when using a partial-least square, we kept the 11 spatial statistics for the parameter estimation.

#### Detection of outlier images

Having validated the inference method, we performed the ABC on the 345 observed landscapes from the global dryland dataset, and we estimated the posterior distribution of *θ* for each landscape (see as an example, posterior densities of two sites in Fig. S26 and comparison between observed and simulated landscapes from the posterior distribution of four sites in Fig. S27). We noticed that the minimal model covered well the observation but lacked of spatial heterogeneity compared to the observed landscapes (see Fig. S28 for comparison of spatial statistics between observations and accepted simulations). Interestingly, for some sites, we found a bimodality in the posterior parameters reflecting the existence of contrasting scales in the image (*e*.*g*., two types of vegetation) (see the example given in Fig. S29). This can for instance be the case in savanna ecosystems, where the coexistence large trees with smaller grasses generates two contrasting scales in the image. The VDmin model does not account for such cases since it models a single class of organism. We therefore chose to remain conservative and systematically detected these landscapes by testing for bimodality in the posterior distribution of parameters. Specifically, we fitted a Gaussian mixture to the posterior distributions of each site, and we removed the sites for which the bimodality of *p* or *q* was significant (*p-value ≤* 0.05 for Hartigan’s dip test; *n* = 52). This reduced the number of observed landscapes to 293. We performed this step using the function *dip*.*test* from the R-package **diptest** (v0.76.0).

Note that our approach coupling parameter inference together with the scale layer could still be applied to such landscapes with two vegetation types provided that a suitable minimal model (*e*.*g*., one with two classes of organisms: grasses and trees for savannas) is identified. In the same vein, in our minimal model, the degraded state is one without vegetation, but the description of other types of transitions such as grass-shrub transitions (e.g. shrub encroachment) might require adopting a different minimal model.

#### Resolution requirement of the inference approach

As previously mentioned, as soon as the image resolution allows the identification of individual plants, our approach performs well thanks to the scale layer, and that, no matter how high the spatial resolution of the image is. When the spatial resolution gets too low, however, the approach may not perform as well because individuals are not well identified, in which case the pixel size might be larger than the size of a plant. We illustrate this resolution limitation by artificially decreasing the spatial resolution of the images in the dryland data set. To do that, we performed a coarse-graining of the observed images using either 2 × 2 or 3 × 3 submatrices as explained in (*37, 38*). Specifically, each submatrix is replaced by its average to obtain a “coarse-grained matrix” with two-times or three-times less pixels in it. Next, for each coarse-grained image *l*, we estimated the posterior distribution of the parameters (*θ*_*l*_), and compared (*i*) the closest selected spatial statistics with the observed ones using the NRMSE computed for each spatial statistic (following the definition given in Eq. S3, S4), as well as (*ii*) the median of the posterior distribution of the parameters for each image and for each spatial resolution. In Fig. S30a, we see that when the spatial resolution decreases, the closest selected simulations become more different compared to the observed landscapes (*i*.*e*., NRMSE increases toward 1). This is especially observed for metrics related to the spatial autocorrelation of the vegetation (Moran I, Spectral ratio, number of neighbors and clustering), and when the spatial resolution gets very low (resolution divided by 3). In fact, in the majority of these 3 × 3 coarse-grained images, the spatial resolution is between 0.4 and 0.6 meters, which blurs the identification of individual plants that typically are of size between 0.25 and 1m^2^ in drylands (*e*.*g*., (*48, 49*)), and also breaks the fine-scale spatial structure of the vegetation. Consequently, when looking at the estimated parameters *p* and *q*, we see that there are important changes in the median of the posterior distribution of *p* and *q* for the landscapes whose spatial resolutions have been divided by three (Fig. S30b). Note that we logically see that the parameter of the scale layer is estimated to be lower compared to when the spatial resolution is higher (Fig. S30b consistently with what has been shown above). **We therefore caution against the use of our approach when the spatial resolution is lower than the typical organism size in the studied system. The lowest spatial resolution on which our approach can be applied is thereby highly dependant on the studied system, but here, in our dryland data set, a threshold of 0.4 meters could be defined**.

### Quantifying the resilience and the vulnerability of sites

#### Bifurcation diagrams

For the 293 retained observed landscapes, we used the posterior distribution of the parameters to predict how a given landscape is expected to change under a sustained increase in the level of abiotic stress. To do so, we assumed that an increase in abiotic stress, such as aridity in our case (*e*.*g*., (*50, 51*)), decreases the parameter linked to plant reproduction (i.e. *p*).

Thereby, for each landscape *l* and sample 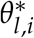 from the posterior distribution *θ*^*∗*^, we decreased 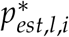 with fixed steps (0.005) until we reached the degradation point, hereafter referred to as the desertification point (*p*_crit,l,i_), which is the maximal value of the parameter *p* for which there is still vegetation. We performed this for each of the 100 pairs of parameters, together giving 100 bifurcation diagrams of the vegetation cover against the parameter *p* per observed landscape *l*.

Note that for high values of *q* the minimal model exhibits discontinuous transitions to degradation with hysteresis (Fig. S14). This means that in this case, the system has two tipping points occurring at different values of *p*: the one at which the system shifts from a vegetated to a desert state (the degradation point, the one of interest here) and the one at which the system would recover from a desert to a vegetated state (the recovery point). Since our approach relies on using the spatial structure of the vegetation to estimate the resilience of the system, it can only be applied to the vegetated state and therefore to the shift from a vegetated to a desert state; indeed, once the system is a desert, there is no spatial structure left in the system that can be used to estimate the distance to the recovery point.

#### Estimating site resilience

We estimated the absolute distance to the degradation point from bifurcation diagram *i* of landscape *l, Dist*_*l,i*_ in Fig. 1, as:

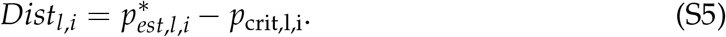

We also computed the relative distance to the degradation point *Dist*_*R,l,i*_ as:

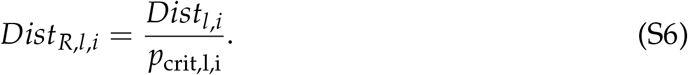

Since we computed these distances to degradation points for each pair of parameters from the posterior (*n* = 100), this gave a sample of 100 distances to the desertification point for each landscape.

### Resilience along gradients of *p* and *q*

We above built the bifurcation diagrams by varying *p*. Our intuition is that *p* is the one that best represents how climate (or a spatially-homogeneous driver) would affect the ecosystem. Interestingly, we observe qualitatively similar fragility of the sites in the dataset when varying *p* or when varying *q* (Pearson correlation = 0.94, p-value < 0.001; n=212), suggesting that our approach is robust when varying either one of the two parameters in isolation (1D). This simplification of real system to an 1D problem (along *p*) does not however capture that multiple drivers can affect the system (e.g., aridity and herbivory), which can translate into changes in both parameters *p* and *q* simultaneously. We tested whether the resilience estimated when decreasing *p* was similar to the one estimated when decreasing *p* and *q* simultaneously (using steps of size 0.0025). Fig. S31a displays the comparison between both estimations, and shows that despite changing both parameters simultaneously, we recover qualitatively similar estimations for the distance to the degradation point of each site (Spearman correlation > 0.98). Since the results when covarying both parameters are similar to those obtained along a *p* gradient (Fig. S31b;or the *q* gradient), they are thus not further discussed here.

### Validating our approach using other models

We validated our approach by comparing the predictions made by the inference with other models generating self-organized patterns and for which the distance to the degradation point can also be computed. We tested two models generating self-organized patterns and possible tipping points to degradation despite different microscopic rules compared to the minimal VDmin model.

First, we used a dryland vegetation model that describes with more details the life-history traits and dynamics of plants in a landscape (*e*.*g*., dispersal, competition, recruitment, mortality; (*16*)). This model allows validating the use of a minimal model by checking whether it replicates well the dynamics of more complex models. In addition, to assess the generality of the approach, we used a spatial model of mussel-beds dynamics (*24*). Importantly, we could have used more models, but because both models belongs to the same types of models (meaning models which, despite relying on different microscopic rules, generate similar types of irregular spatial structure (*25, 40, 52*)), we restricted our analyses to these two. We give more information about these two models below (see Fig. 2 for model illustration).

#### Model of dryland vegetation dynamics with life-history traits

The first model is a spatial model of vegetation dynamics in drylands that has seven parameters describing the life history of plants in arid ecosystems ((*16*) ; Fig. 2). Each site in the landscape can be in one of three states: degraded (*D*), empty but fertile (*F*), or colonized by vegetation (*V*). Vegetation produces seeds that are dispersed both locally in the four nearest-neighbors (with a fraction *δ*) and across the whole landscape (with a fraction 1 − *δ*). Seeds can only germinate on a fertile site. Let *ρ*_*V*_ the fraction of sites covered by vegetation and *q*_*V*|*F*_ the local fraction of vegetated sites in the neighborhood of a fertile site (4 nearest neighbors), then the probability of seed recruitment is given by:

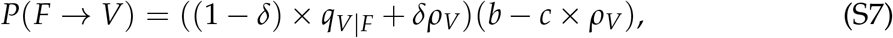

where *b* is the recruitment probability of seeds, and *c* is the density-dependent competition for water. Once colonized a fertile site, the vegetation can die with a mortality rate *m*

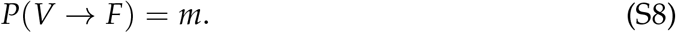

The regeneration of a degraded site occurs at a basal rate *r* and is increased if neighboring sites are vegetated (*i*.*e*., facilitation):

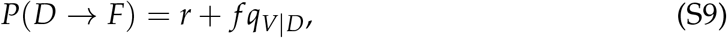

where *f* is the strength of facilitation, and *q*_*V*|*D*_ is the local fraction of vegetated sites in the neighborhood of a degraded site. Last, the degradation of a fertile site occurs at a rate *d*:

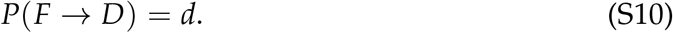

We ran simulations on different parameter sets. To capture different spatial structure and vegetation cover, we varied the strength of facilitation (*f* ∈ {0.5, 0.75, 0.9}) and the recruitment probability *b* (20 values equally spaced between 0.4 and 0.9), and fixed other parameters (*r* = 0.11, *d* = 0.1, *m* = 0.1, *δ* = 0.1, *c* = 0.1 following (*16*)). The simulations were performed similarly to the main VDmin model with a 1500 time step burning period before computing the spatial statistics. We applied the inference approach on these 60 “virtual landscapes”, estimated their parameter posterior distributions *θ*^*∗*^, and computed the distance to the degradation point. To compare with, we computed the distance to the degradation point in the Kéfi model, assuming that an increase in aridity would decrease the recruitment rate of seedlings (*b*).

#### Model of mussel-beds dynamics

Besides drylands, other systems such as mussel-beds exhibit self-organized patterns characterized by a power-law distribution of patch sizes and for which, spatial structure of mussels is a good indicator of the resilience of the system (*15, 24*). To emphasize the generality of our method, we used a second model developed by Guichard et al. (*24*). In this model, each site in the landscape can be in one of three states: empty (*E*), colonized by mussels (*M*), or disturbed (*D*) (Fig. 2). Colonization of an empty site by mussels occurs at a rate proportional to the recruitment rate *α*_2_.

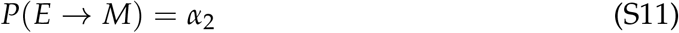

A mussel can become disturbed at a rate basal rate *δ*_0_, that is increased by a density-dependent disturbance rate *α*_0_ when a mussel has at least one disturbed site in its neighborhood (Fig. 2):

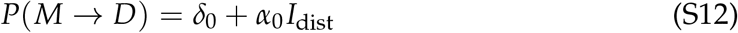

with *I*_dist_ = 1 if there is at least one disturbed site in the neighborhood of a mussel site, and *I*_dist_ = 0 if not. After one time-step, each disturbed site is automatically stabilized to an empty state (*d* = 1).

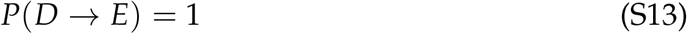

We ran simulations with 10 values of *δ*_0_ equally spaced between 0.01 and 0.3, three values of *α*_0_(∈ {0.1, 0.2, 0.3}), and two of *α*_2_(∈ {0.6, 0.8}) to generate virtual landscapes with different mussel cover. Next, we used our inference approach on these “virtual landscapes”, estimated the posterior distributions *θ*^*∗*^, and computed the distance to the degradation point. To compare with, we computed the distance to the degradation point in the Guichard model, assuming that an increase in environmental stress would increase the disturbance rate (*δ*_0_).

For both the mussel-bed and the dryland vegetation models, we quantified whether our inference approach could accurately rank sites from the closest to the farthest of their degradation point using the Spearman rank correlation. To account for the posterior predictive uncertainty, we sampled 500 times each inferred distance to the degradation point, assuming that the distance to the degradation point of each simulated landscape followed a normal distribution with mean and standard-deviation computed on the distribution of 100 distances to the degradation point.

### Assessing the drivers of global drylands resilience

Because our approach only relies on a snapshot of a site, it does not account for the effect of variability at relatively short time scale such as seasonality. What we detect on those images are essentially perennial plants and their spatial structure can be considered to be a slow variable. In fact, spatial structure hardly changes within the year (annuals generally hardly affect the spatial structure of drylands), and the year-to-year spatial structure changes reflect mortality of adult individuals (which again are slow processes since a shrub species can live between 20 and 200 years, see review (*53–55*), and recruitment of new individuals are rare events in drylands. The very few studies that have investigated temporal changes in spatial vegetation structure have shown that these changes are quite slow: Deblauwe et al (2012) showed that upslope migration of perennials was a very slow process (slower than 1 meter per year; (*56*)). We therefore believe that seasonability does not significantly affect the resilience and statistical results presented.

#### Proximate drivers

We tested how the vegetation cover and the spatial structure of the vegetation were related to the estimated distance to the desertification point. To do that, we fitted a mixed-effect model with the median of the distribution of the distance to the desertification point as a response variable, and both the spatial autocorrelation of the vegetation (Moran I) and the vegetation cover as predictors. The site was included as a random intercept. This statistical model with the two predictors was then compared to statistical models with only one of the predictors (either the vegetation cover or the spatial autocorrelation of vegetation alone) using the bootstrapped-AIC of each model.

#### Ultimate drivers

Next, to assess the effects of the soil, climatic and vegetation attributes on drylands resilience, we divided our analyses in two parts. First, we used mixed-effects models to investigate how each predictor was related with the distance to the desertification point. Then, we explored the indirect effects of each driver on the distance to the desertification point.

To assess the drivers of drylands resilience, we fitted a linear mixed-effect model with the estimated distance to the desertification point (*Dist*) as the response variable. Let *Dist*_*l,s*_ be the median of the posterior distribution of the distance to the desertification point of landscape *l* in site *s*, the model reads as follows:

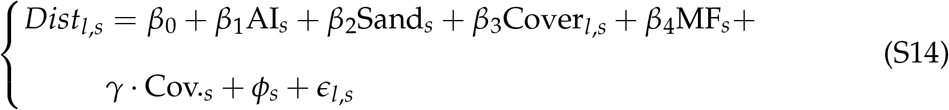

where *β*_0_ is the global intercept, AI is the aridity level, Sand is the soil sand content, and MF is the multifunctionality index. We also included geographical variables (Cov.) to limit potential confounding effects; namely, the latitude, longitude (cos and sin-transformed), the slope and the elevation. The predictors were weakly correlated (see Fig. S9 for the pair correlations). The site was included as a random intercept (*ϕ*_*s*_).

Before running the models, we examined the distributions of all drivers, and tested their normality. We log-transformed the distance to the desertification point to improve normality. All variables were centered and reduced. We also inspected the distribution of residuals and the model assumptions using the *mcp*.*fnc* function from the package **LMERConvenienceFunctions**. Last, we tested for model multicollinearity using Variance Inflation Factors, and tested the potential spatial autocorrelation in the residuals using Moran tests, but neither multicollinearity nor spatial autocorrelation were observed in the model (Table S1).

From the fitted model, we then used model partial residuals to show how each predictor influenced the distance to the desertification point (Fig. 3A-C). Also, because we only had facilitation data for 184 among the 293 landscapes (70 over 115 sites), we performed the same models while including the level of facilitation in the predictors in a separate analysis.

Fitting the model using the relative distance to the desertification point led to qualitatively similar results (Fig. S32).

All models were fitted using the median of the posterior distributions of *q*_*est*_ and the distance to the degradation point. We used the *lmer* function from the **lme4** package.

#### Indirect effects using SEMs

Next, we plugged all drivers together using Structural Equation Modeling (SEM) to investigate the indirect effects of soil, climate and vegetation attributes on the estimated distance to the desertification point. A SEM complements the mixed-effects models by partitioning the influences between multiple variables, and separating the indirect effects of the drivers on the distance to the desertification point. The first step of a SEM analysis is to establish an a priori model based on the known effects and relationships among the drivers of the distance to the desertification point. We designed a SEM where the distance to the desertification point is influenced by proximal drivers (the vegetation cover and the spatial autocorrelation of vegetation, Moran I) based on results of previous paragraphs, that are themselves modulated by ultimate drivers (multifunctionality, aridity). We tested for a link between aridity and multifunctionality following (*1*). We did not include facilitation in the SEM given that it was measured on a subset of all data (*n* = 184).

Before running the SEM, we examined the distributions of all drivers, log-transformed the distance to the desertification point, and centered and reduced all variables. In addition, we controlled for potential effects of geographical variables not included in the SEM by fitting the SEM on the residuals of a linear model with the distance to the desertification point as response variable and the latitude, longitude (cos and sin-transformed), the slope and the elevation as predictors.

We tested the goodness of fit of our model using the Fisher-C statistic. The model has a good fit when the associated p-value is high (*e*.*g*., by convention > 0.05), which is the case here (Fig. 3). Last, we calculated the standardized total effects of aridity and multifunctionality on the distance to the desertification point by multiplying links along the paths. All coefficients were evaluated by bootstrapped (1000 replicates; see Table S2 for the confidence intervals on path coefficients). SEM analyses were performed using the **piecewiseSEM** and the **semEff** packages. The SEM were done with the absolute distance to the desertification point, but similar results were obtained using the relative distance (results not shown).

### Identifying vulnerable sites

Drylands are generally experiencing an increase in aridity levels (*50*), which consequently drives the desertification of large areas (*51*). To turn our approach into a tool to identify the most vulnerable dryland sites of our dataset, we related the log-transformed estimated distance to the desertification point with the predicted change in aridity of each site in the next century (from 1950 to 2100) under the IPCC’ RCP 8.5 scenario that assumes a sustained increase in carbon emission from human activities. We obtained these climatic data from the European Copernicus climate change service (*57*). We derived the changes in aridity (measured as annual evaporation divided by precipitation, monthly averaged) of each site as the mean slope of aridity change between 1950 and 2100. Since different CMIP5 models were available (*n* = 9), we average the slope obtained from the different climatic models. It is noteworthy that sites experiencing currently relatively low aridity levels, are projected to undergo faster increase in aridity in the next decades (Fig. S10). Then, we defined the five vulnerability-based areas in Fig. 4 using a K-means clustering approach on the space defined by the distance to the desertification point and the projected aridity change.

Note that the use of several different criteria for the selection of the number of clusters did not lead to a clear consensus (preliminary results not shown). We therefore chose 5 clusters for their interpretability on the 2D-space, and we only used them for graphical representation purposes.

## Acknowledgments

We thank Fernando T. Maestre for sharing the BIOCOM dryland dataset. We also thank Ismaël Lajaaiti, Jean-François Arnoldi and Nathan Humbert for feedbacks and as well as the BioDICée team discussions on the project.

## Author contributions

Conceptualization: SK, BP; Methodology: SK, BP, SD; Investigation: BP; Visualization: BP, SD, IG, SK; Funding acquisition: SK, BP, SD, IG; Supervision: SK, SD, IG; Writing – original draft: BP; Writing – review and editing: BP, SD, IG, SK.

## Data and materials availability

The data and the code for reproducing results are available through Zenodo (*48*).

