## Supplementary_figures_text for "Estimating distances to desertification points from dryland ecosystem images"

### Supplementary text

##### Assessing the robustness of the minimal model

We evaluated the ability of the minimal model to *(i)* generate similar spatial patterns as the observed ones, *(ii)* provide a reliable ranking of sites based on their vulnerability to increasing aridity, and *(iii)* encapsulate many parameters of more complex models into two parameters $(p, q)$.

First, by comparing each spatial statistic computed on the simulated landscapes from the two complex models with the selected ones from the inference approach, we show that each component of the spatial structure of the simulated landscapes are well covered by the minimal model (Figs. S33-34). Despite its minimalistic rules with two parameters, the minimal model therefore allows covering the diversity of spatial patterns generated by more complex models exhibiting spatially irregular spatial patterns. This also means that the three models we used in this study belong to the same types of spatial models generating similar kinds of irregular patterns. In addition, we show that, while the distance to the degradation point (distance to a desert/no-mussel state) quantitatively change depending on the model used (Fig. 2), the estimated distance to the degradation point is qualitatively similar across models, meaning that two sites with different spatial structure will be ranked in the same order depending on their spatial structure. This also implies that the relative resilience of different ecosystems is consistent across models used, and can be reliably estimated using our inverse modelling approach. The reason for this, is because there is a topological equivalence of the bifurcation diagrams across the models showing similar irregular spatial patterns, making the distance to the degradation point qualitatively consistent across models despite their diversity of rules.

It is noteworthy that while our inference approach, boiling down the spatial structure of ecosystems to two parameters, allows quantitatively ranking sites based on the distance to their degradation point (Fig. 2), it cannot predict well the type of transition (abrupt or gradual) because the latter is a property of the model rather than a property of the emergent spatial structure of the vegetation. In fact, preliminary analyses using the mussel-bed and the dryland vegetation model showed that the loss of vegetatation or mussel density at the degradation point could not be well estimated using the minimal model. Since the type of transition cannot be predicted from ecosystem images (it is not a property of the spatial structure of the system but is rather model-specific), we therefore focus on the distance to a degradation point.

On a final note, we observe that the estimated distance to the degradation point changes in an expected way along model parameters of the dryland and mussel-bed models (Fig. S35). For instance, the estimated distance to the degradation point linearly increases with the recruitment of vegetation ($b$) or the facilitation parameter ($f$) in the vegetation model. In the mussel-bed model, the estimated distance to the degradation point linearly decreases with the mortality rate of mussels ($d$) while increasing with the colonization rate of mussel ($a_2$). Together, we show that the minimal model allows encapsulating the dynamics and the spatial structure of more complex models, estimating well the relative vulnerability of the ecosystems, but not predicting well the type of transition. In other words, assuming that the observed ecosystem can be described by the same minimal model, we show that the approach allows estimating a minimal and informative parametrization of the dynamical state of the observed ecosystem.

### Supplementary figures (S1-S35)

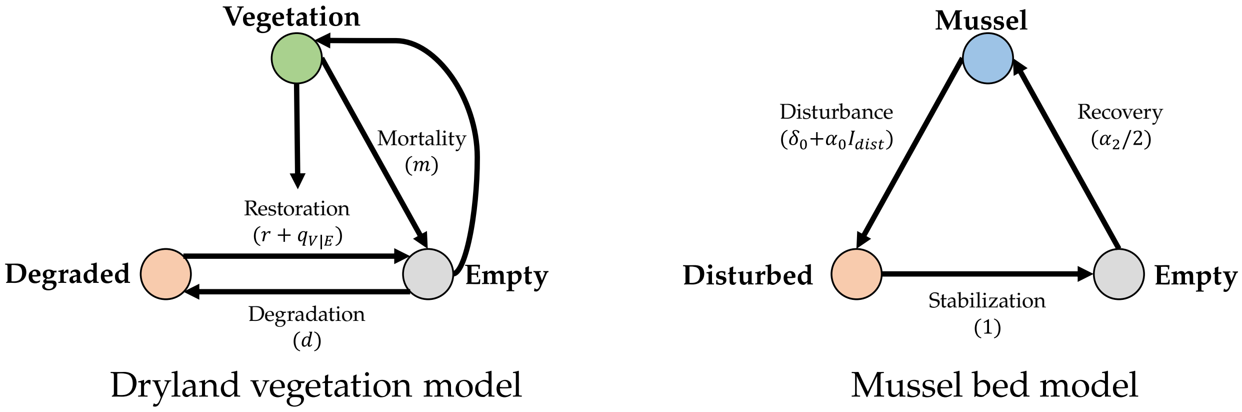


Fig. S1: **Structure of the two models used for the validation of our approach**.
We used two models to validate our approach. The first (left), of Kéfi et al. (*16*), is a model of dryland vegetation dynamics that includes detailed life-history traits (dispersal, recruitment, restoration by facilitation). The second model is a mussel-bed model of Guichard et al. (*24*) and describes the dynamics of disturbances in mussel-beds in intertidal ecosystems. For each model, we show the three different states as well as the probabilities of transitions between each states. The model parameters are described in Methods.

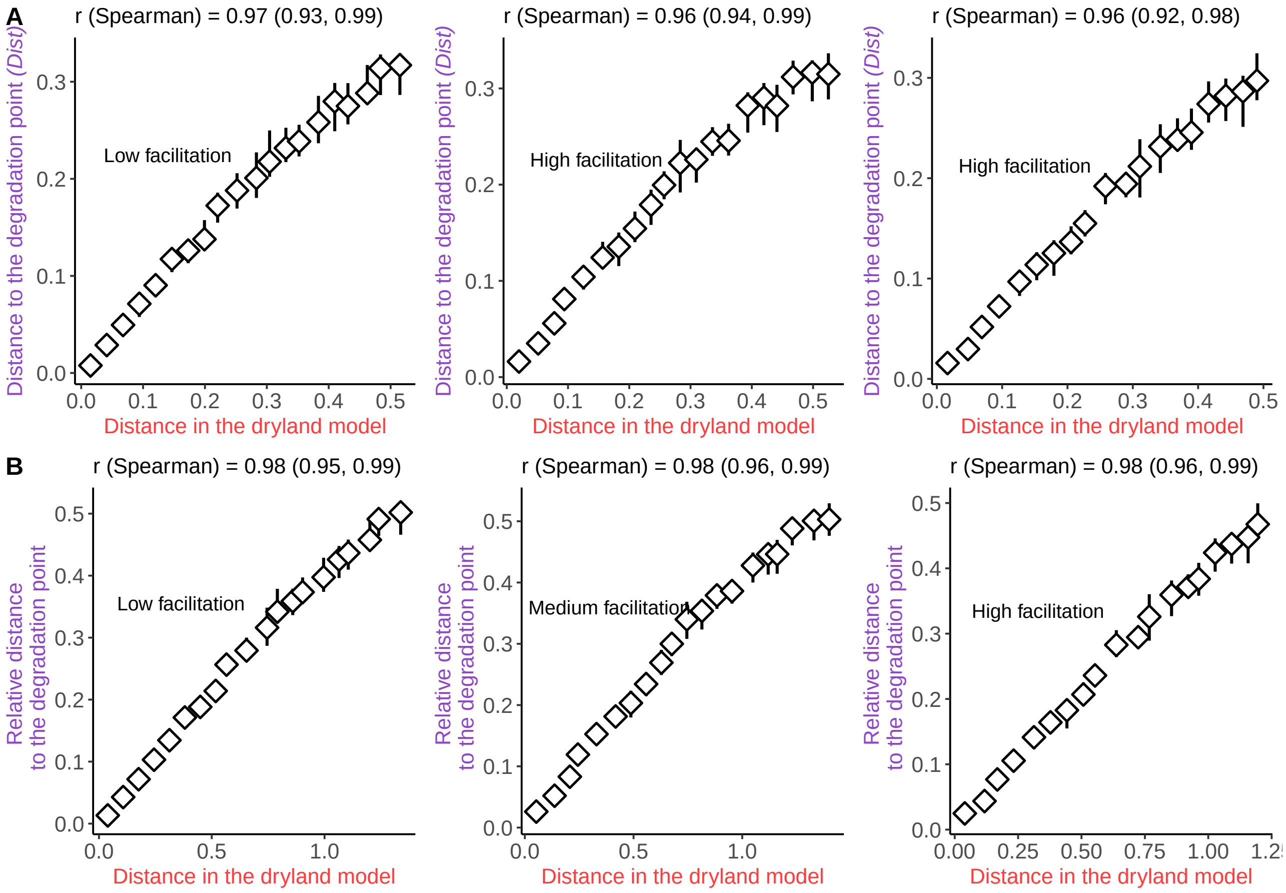


Fig. S2: **Our approach performs well at ranking simulated vegetation landscapes based on their distance to the desertification state.**
We used the dryland vegetation model of Kéfi et al. (*16*) with different sets of parameters. For each parameter set, we estimated the "true" theoretical distance to the desertification point by decreasing the probability of recruitment of plants (i.e. the parameter $b$; x-axis). Then, we compared this theoretical prediction with the inferred distance to desertification point using the inferred posterior distribution of parameters $p_{est}$ and $q_{est}$ of each simulated landscape from the minimal model. Points and their associated range represent the median surrounded by the first and third quartiles of these distributions.
(A) Absolute distance to the degradation point ($p_{est} - p_c$). (B) Relative distance to the degradation point ($(p_{est} - p_c)/p_c$).
The median ($r$) of the Spearman correlation as well as the quantiles at 5 and 95% (brackets) are indicated.

**Fig. S3: Our approach performs well at ranking mussel-bed landscapes based on their distance to the degradation point to the no-mussel state.**

We used the mussel-bed model of Guichard et al. (*24*) with different set of parameters. For each parameter set, we then compared the "true" theoretical distance to the no-mussel state by decreasing parameter $\delta_0$ (x-axis). Then, we compared this theoretical prediction with the estimated distance to the degradation state using the inferred posterior distribution of parameters $p_{est}$ and $q_{est}$ of each simulated landscape. Points and their associated range represent the median surrounded by the first and third quartiles of these distributions.

(A) Absolute distance to the degradation point ($p_{est} - p_c$). (B) Relative distance to the degradation point ($(p_{est} - p_c)/p_c$).

The median ($r$) of the Spearman correlation as well as the quantiles at 5 and 95% (brackets) are indicated.

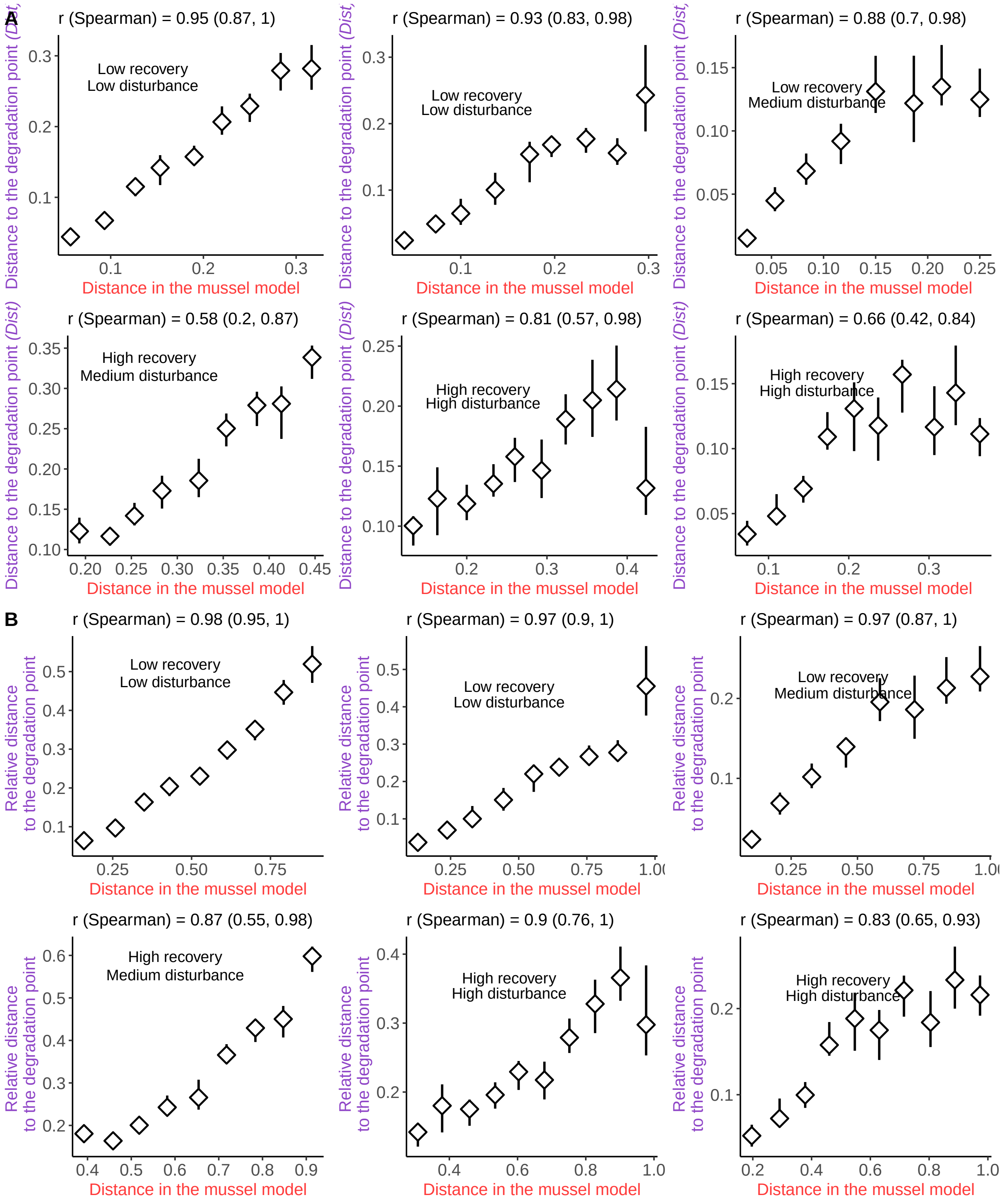
A
r (Spearman) = 0.95 (0.87, 1)
Low recovery
Low disturbance
r (Spearman) = 0.93 (0.83, 0.98)
Low recovery
Low disturbance
r (Spearman) = 0.88 (0.7, 0.98)
Low recovery
Medium disturbance
r (Spearman) = 0.58 (0.2, 0.87)
High recovery
Medium disturbance
r (Spearman) = 0.81 (0.57, 0.98)
High recovery
High disturbance
r (Spearman) = 0.66 (0.42, 0.84)
High recovery
High disturbance
Distance to the degradation point (Dist)
Distance in the mussel model
B
r (Spearman) = 0.98 (0.95, 1)
Low recovery
Low disturbance
r (Spearman) = 0.97 (0.9, 1)
Low recovery
Low disturbance
r (Spearman) = 0.97 (0.87, 1)
Low recovery
Medium disturbance
r (Spearman) = 0.87 (0.55, 0.98)
High recovery
Medium disturbance
r (Spearman) = 0.9 (0.76, 1)
High recovery
High disturbance
r (Spearman) = 0.83 (0.65, 0.93)
High recovery
High disturbance
Relative distance to the degradation point
Distance in the mussel model

Fig. S3:

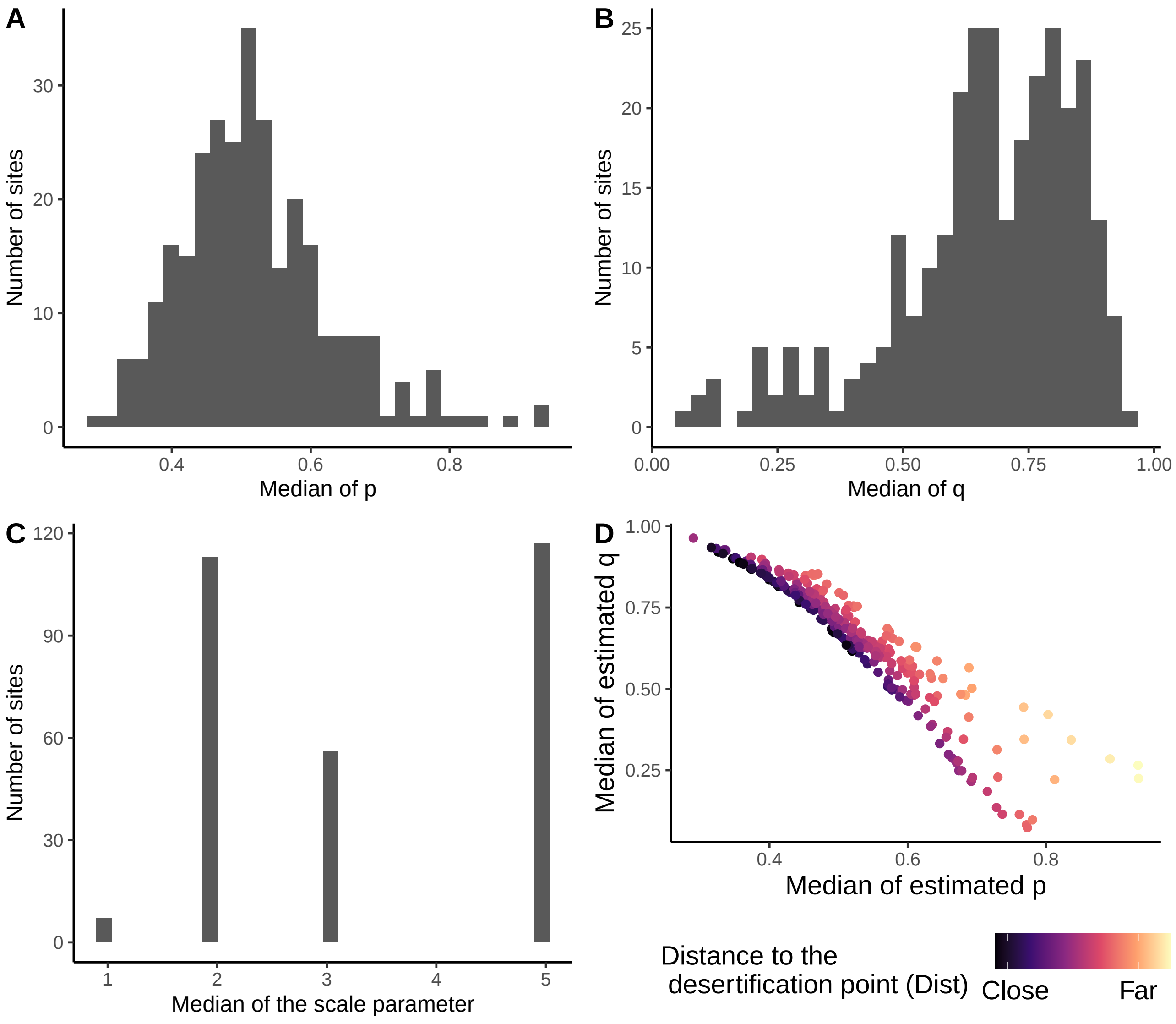


Fig. S4: **Distribution of the median posterior of $p_{est}$ and $q_{est}$ across the 293 sites.** The distribution of the median of posteriors of the 293 sites of parameters $p_{est}$ (A), $q_{est}$ (B), $\eta$ (C) and the joint distribution of $(p_{est}, q_{est})$ (D) are displayed. The color in D represent the distance to the desertification point (*Dist*).

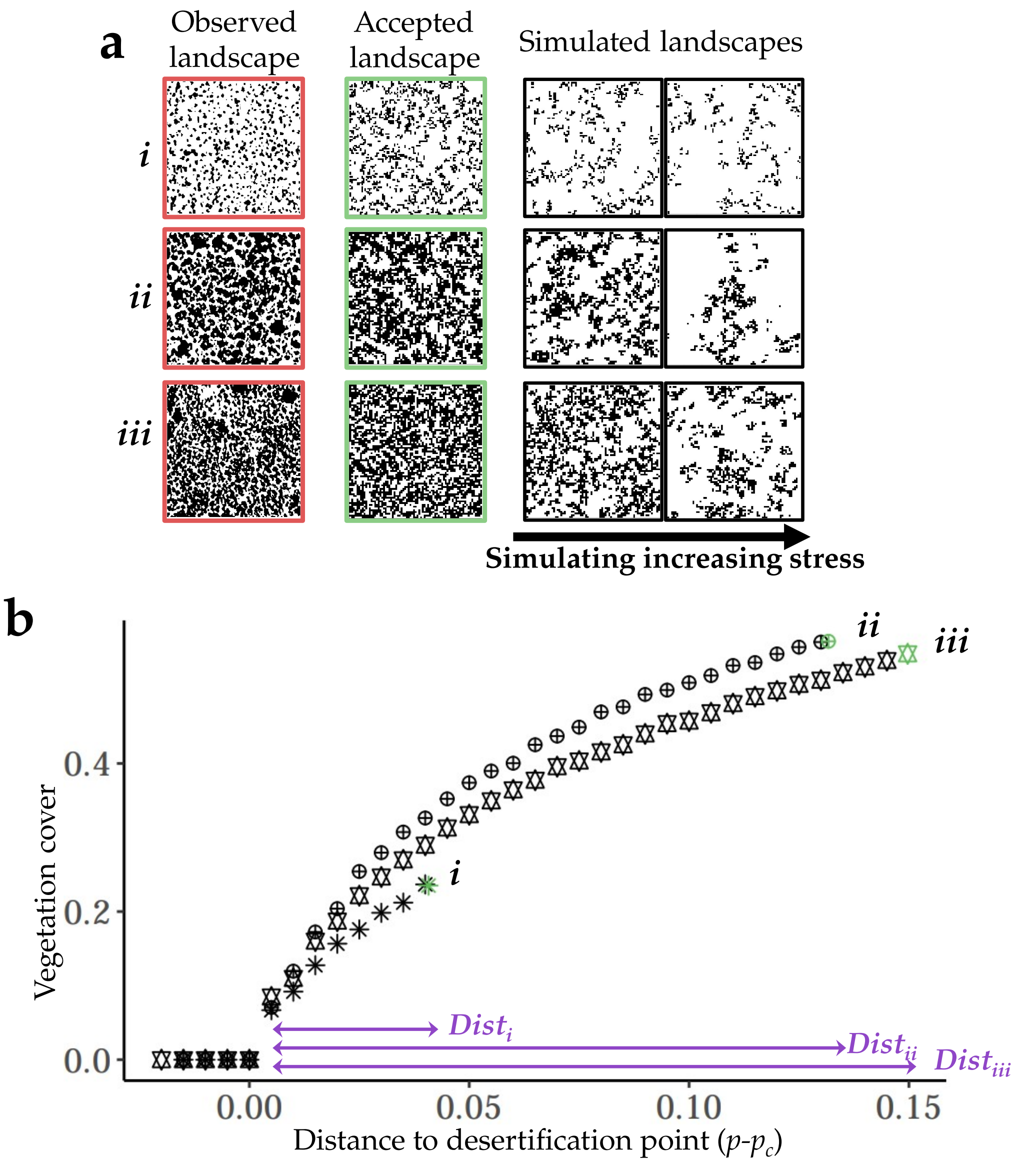


Fig. S5: **A case study of the inference approach on three field sites.**
(A) For each observed vegetation landscape (*i*-*iii*), we performed the ABC inference and estimated the posterior parameter distributions of $p_{est}$ and $q_{est}$. We show an example of a landscape generated using the posterior parameter distribution of each site. We then simulated an increasing stress (decreasing $p_{est}$) until the degradation point is reached ($p_{est} - p_c = 0$). We show the bifurcation diagram for the median of the posterior distribution $q_{est}$. (B) Vegetation cover of each site against the parameter $p_{est}$. x-axis is scaled such that 0 corresponds to the degradation point.

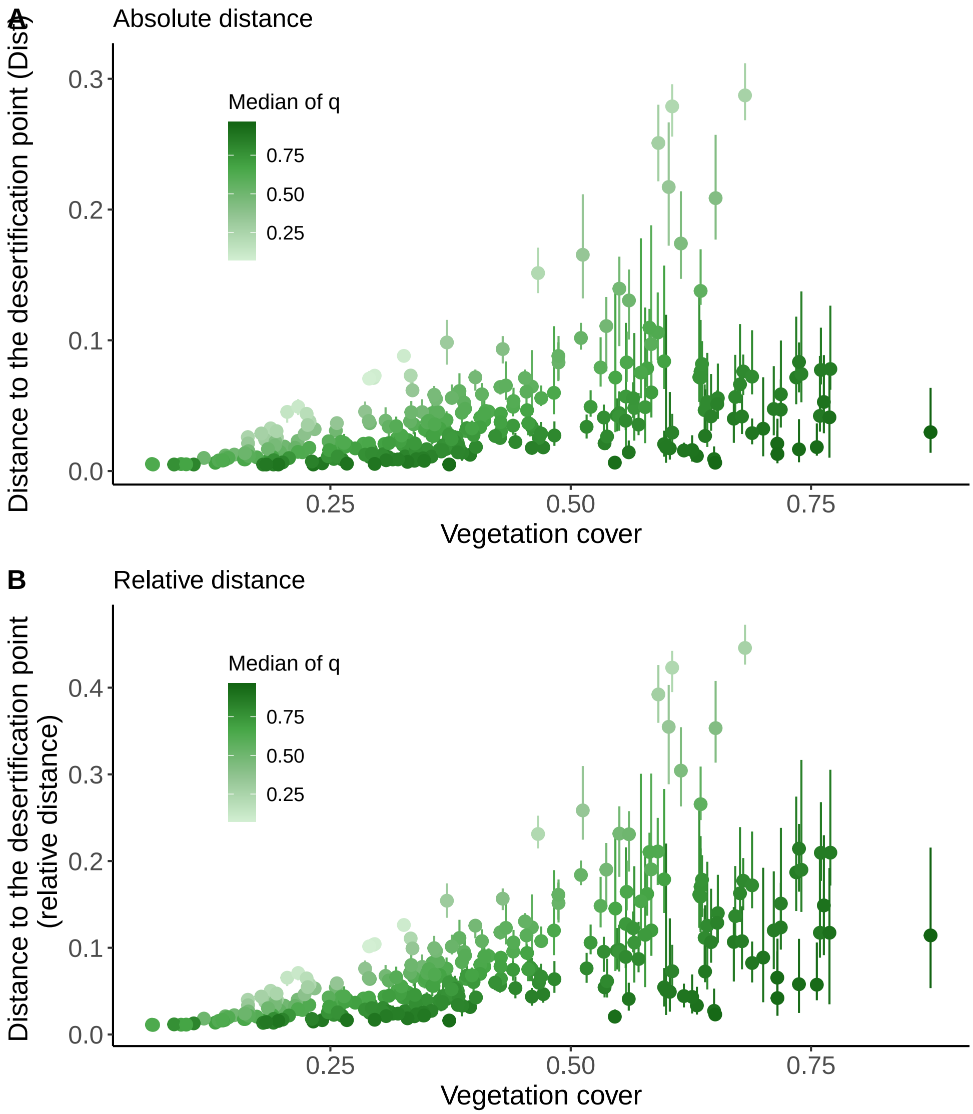


Fig. S6: **Correlation between landscape vegetation cover, estimated parameter $q_{est}$ and the distance to the desertification point.**
Relationship between the estimated distance to the degradation point and the vegetation cover. The color corresponds to the median of the posterior of parameter $q_{est}$. (A) Absolute distance to the desertification point ($p_{est} - p_c$). (B) Relative distance to the desertification point ($\frac{p_{est}-p_c}{p_c}$). While higher vegetation cover and low parameter $q$ seem to be associated with higher distance to the desertification point, alone each predictor is not sufficient to explain the distance to the desertification point.

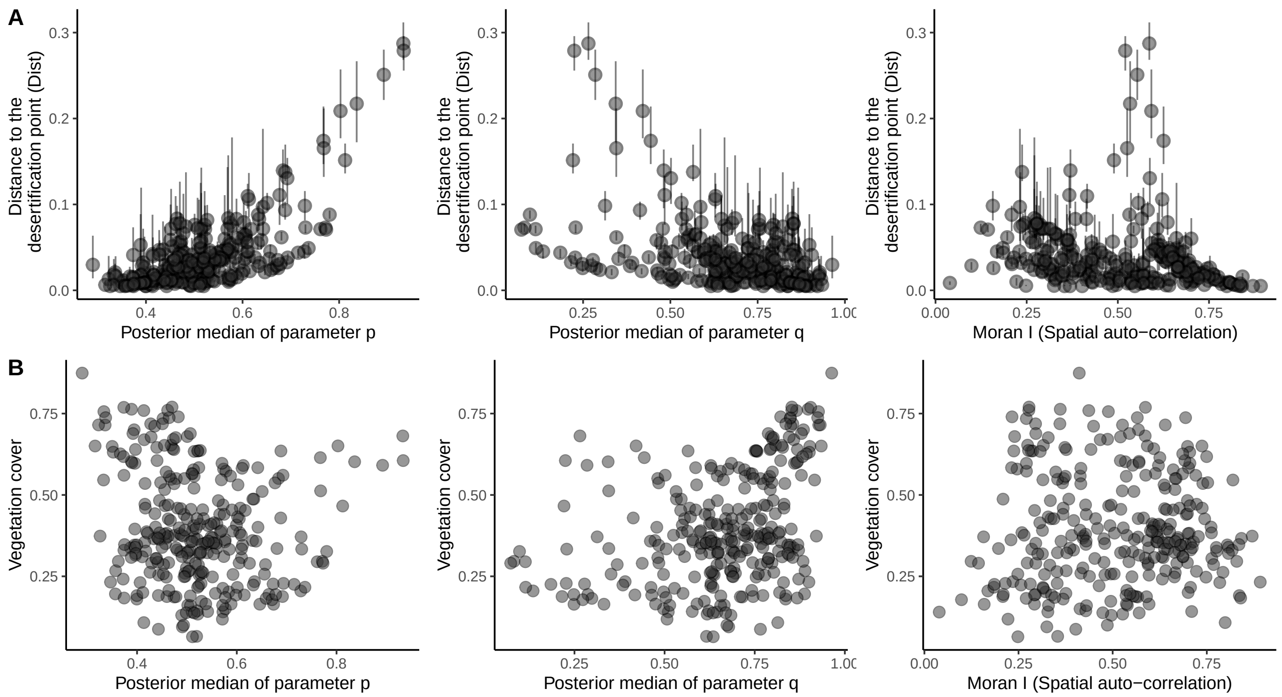


Fig. S7: **Correlation between estimated parameters $p_{est}$, $q_{est}$ and the distance to the desertification point.**
(A) Correlation between estimated distance to the desertification point, the median of the estimated posterior distribution of $p_{est}$, $q_{est}$, and, the spatial autocorrelation (Moran I). (B) Correlation between the vegetation cover of the landscape, the median of the estimated posterior distribution of $p_{est}$, $q_{est}$, and, the spatial autocorrelation (Moran I). Each point is surrounded by the posterior predictive uncertainty (quantiles at 25 and 75%).

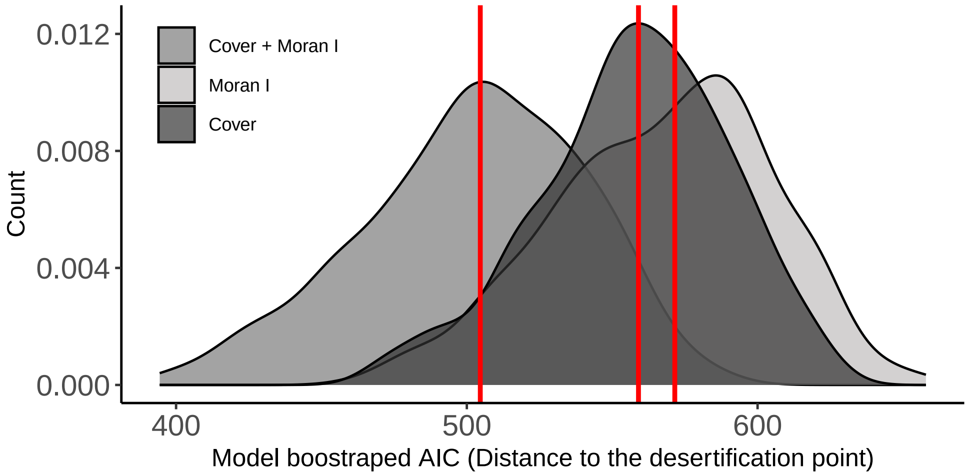


Fig. S8: **AIC comparison reveals that neither spatial structure (spatial autocorrelation, Moran I) nor vegetation cover alone explains the distance to the desertification point: both complements each other.**
Distributions of AIC of mixed effects models using either the spatial autocorrelation (Moran I), the vegetation cover or both as predictors of the distance to the desertification point. The site was included as a random effect. AIC were bootstrapped to obtain these distributions. Red lines correspond to the median AIC.

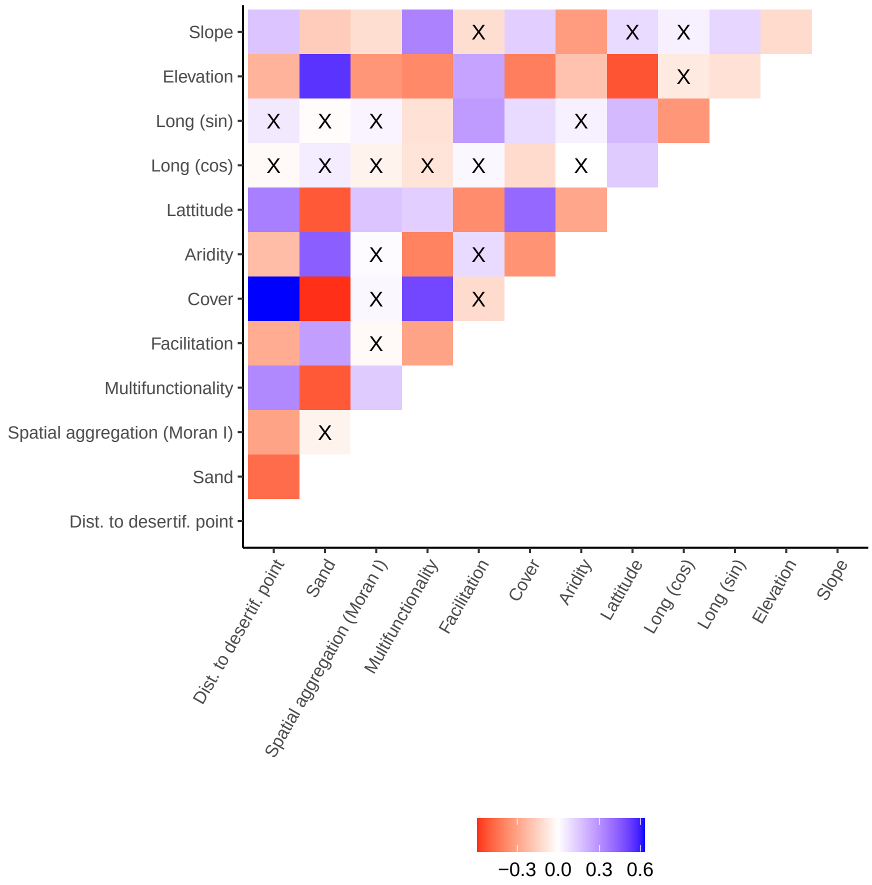


Fig. S9: **Correlation between drivers, estimated parameters and distance to the desertification point.** A black cross indicates non-significant correlation (*p-value* $> 0.05$). Correlation with facilitation were only computed for a subset of $n = 184$ landscapes. The slope corresponds to the measured slope in the field, the longitude was sin- and cos- transformed, aridity is the aridity level, cover is the vegetation cover of the landscape, multifunctionality is the soil multifunctionality as defined in Methods, sand is the soil sand content, and Dist. to desertif. point is the estimated distance to the desertification point (*Dist*).

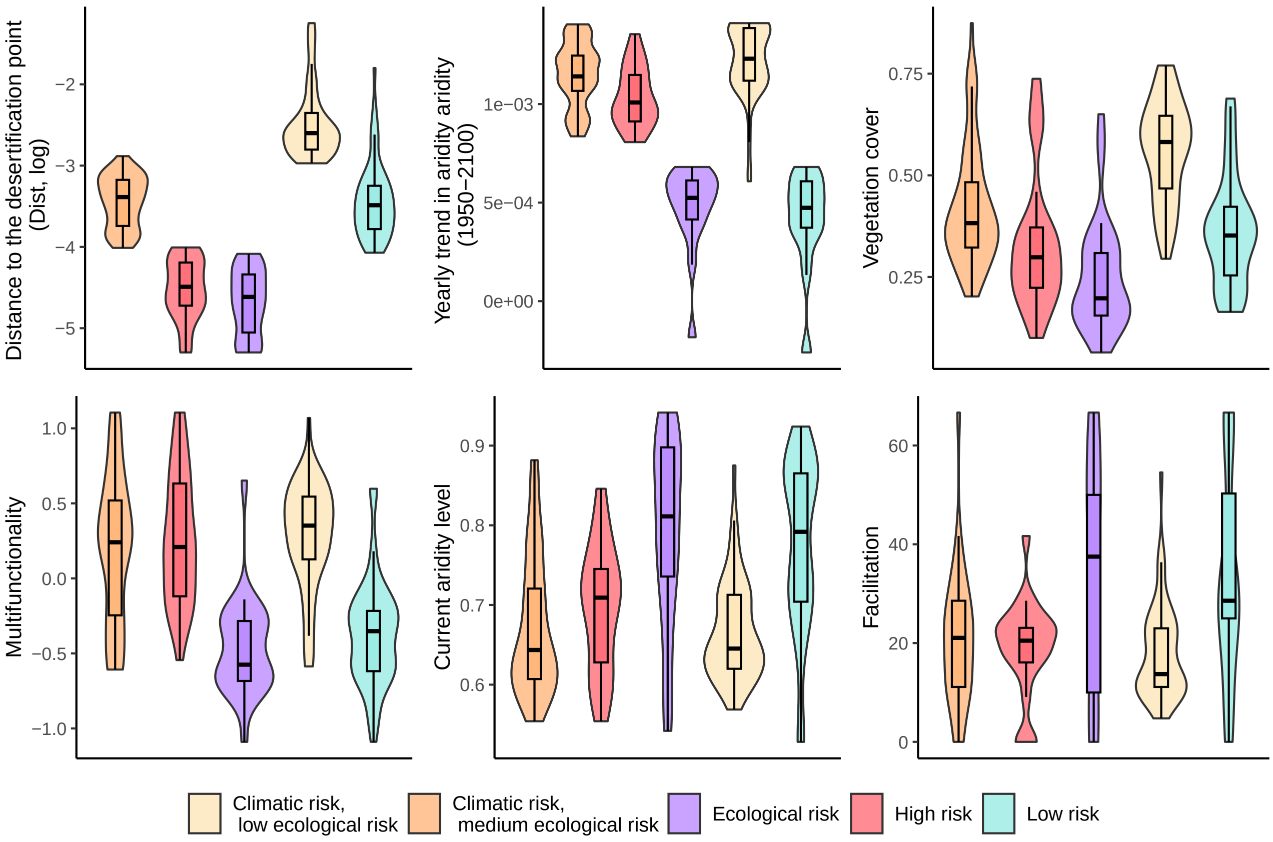


Fig. S10: **Characteristics of vulnerability clusters defined in Fig. 4.**
Distributions of the estimated distance to the desertification point, projected aridity change (annual slope 1950-2100), vegetation cover, current aridity level, soil multi-functionality index, and facilitation strength for the five clusters defined in Fig. 4.

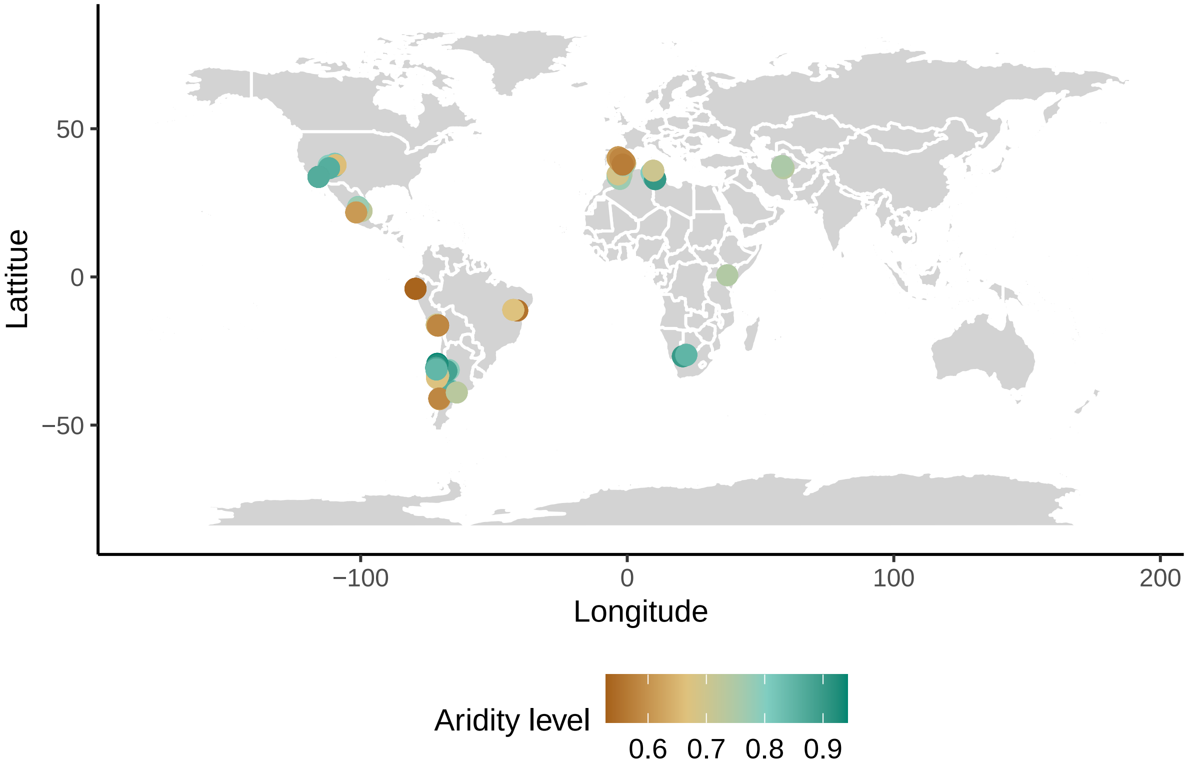


Fig. S11: **Distribution of sampled sites across the globe.**
115 sites in 13 countries were sampled across the globe in different biotic and abiotic (here aridity) conditions. For each site, three images of vegetation landscapes were extracted by Berdugo et al., (2017) (see (*1*, *2*)).

Caption for Fig. S12 **Principal component analysis on the spatial structure of simulated landscapes.**

We performed a principal component analysis using the different spatial statistics computed on a sample of 20000 simulated landscapes from the minimal VDmin model. (A) Colored by the parameter $p$. (B) Colored by the parameter $q$. (C) Colored by the contribution of each spatial statistic to each of the different principal components (PC1 in the left panel, PC2 in the middle panel, and PC3 in the right panel). The combination of the three axis is shown. Cover = vegetation cover in the landscape, # neighbors = mean number of vegetated neighbors in the neighborhood of a vegetated site, Clustering = clustering of vegetation in patches, Skewness = the spatial skewness of vegetation, Variance = the spatial variance of vegetation, Autocorrelation = Spatial autocorrelation of the vegetation (Moran I), SDR = Spectral density ratio, PLR = power-law range, Exponent p.l. = exponent of the fitted power-law, Frac. max = Fraction of the landscape covered by the largest patch, CV PSD = coefficient of variation of the patch size distribution. See Methods for the definition of the spatial statistics used ("Choice of the discrepancy between observations and simulations")

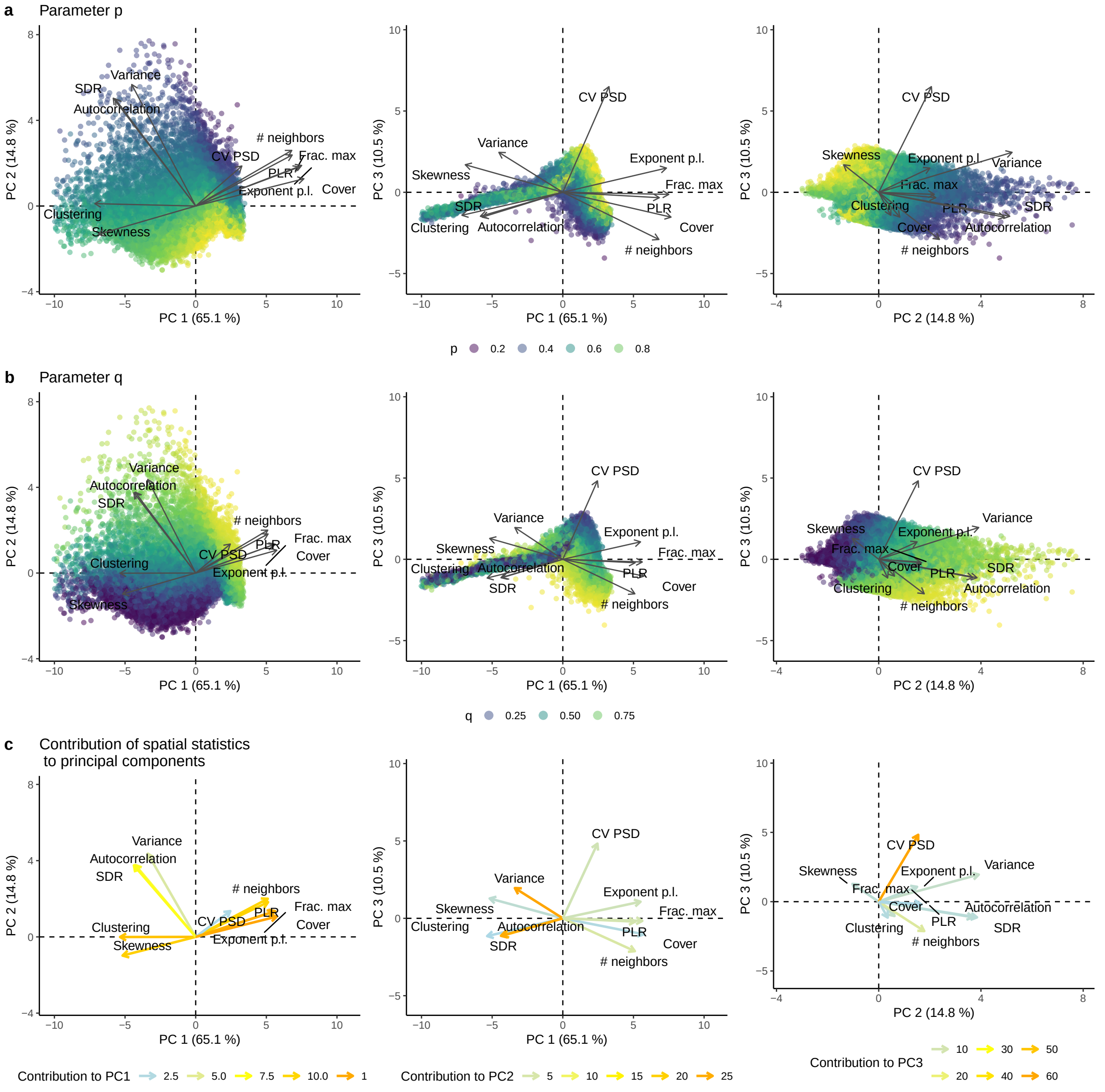


Fig. S12: **Principal component analysis on the spatial structure of simulated landscapes.**

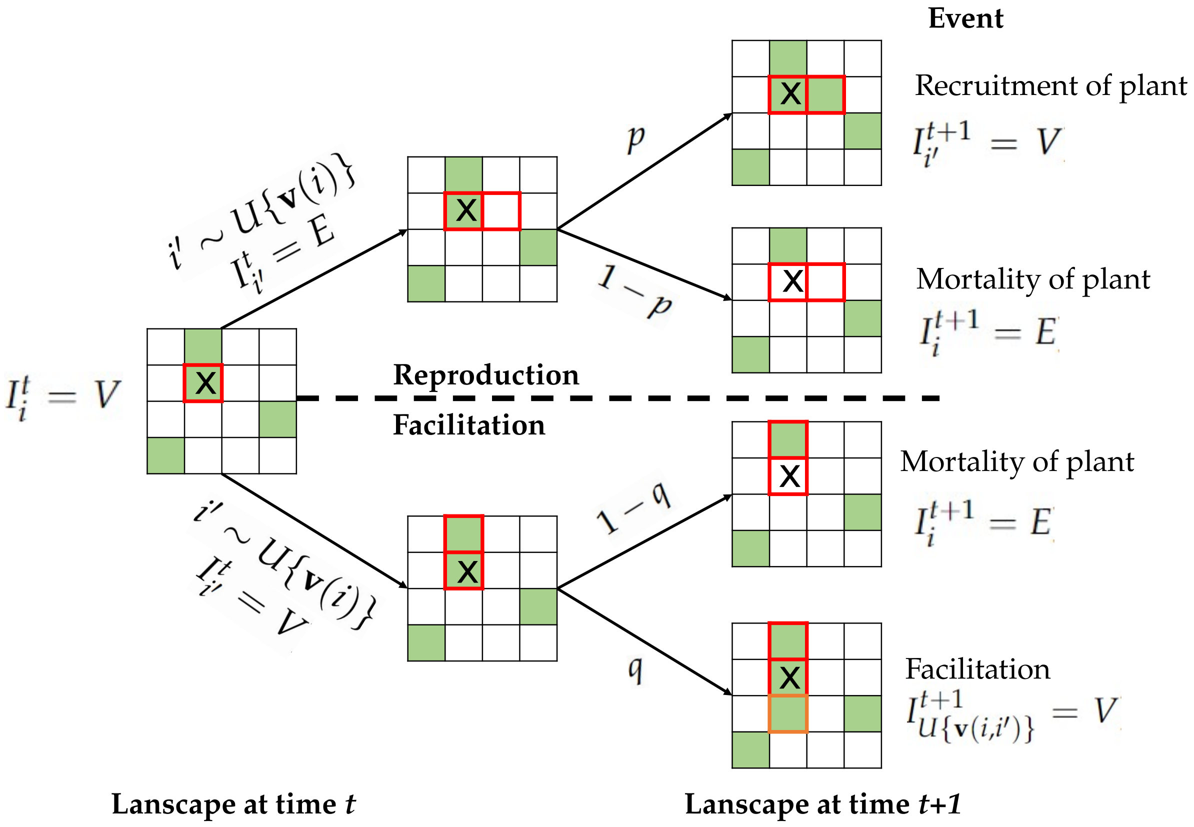


Fig. S13: **Description of the model used.**
Details about the different steps are given in method section. The focal site (with the cross) and its neighbor are contoured in red. Newly colonized site by facilitation is colored in orange.

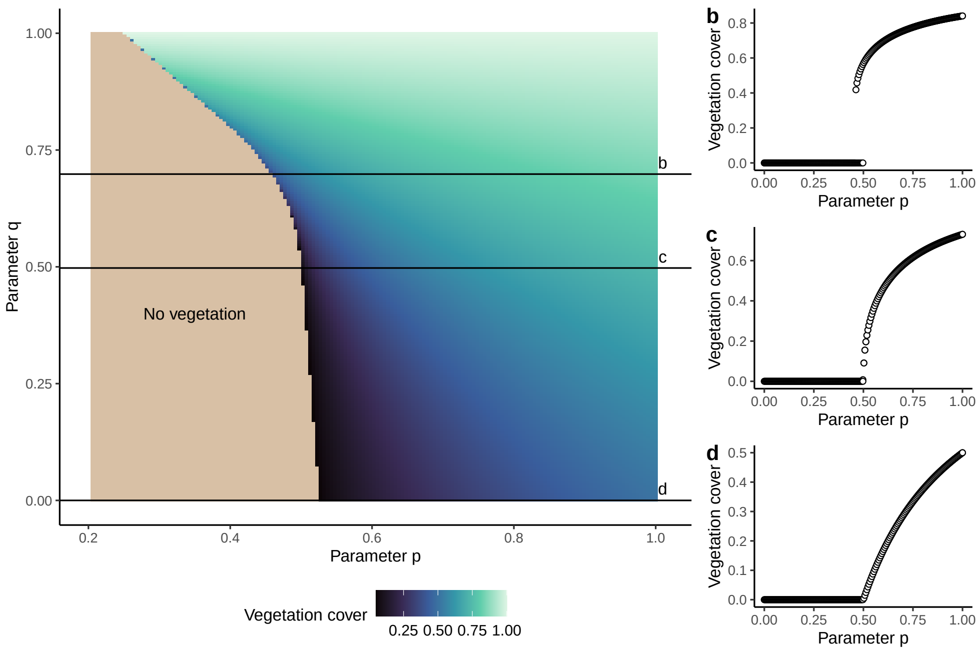


Fig. S14: **The mean-field phase-diagram of the VDmin model.** (a) We solved the mean-field approximation of the model (Eq.S1) for 200 by 200 combination of parameter $p$ and $q$ values, and show the vegetation cover at equilibrium. (b-c) For three transects of parameter $q$, we show the bifurcation diagram (vegetation cover against parameter $p$ value). The model predicts gradual (continuous) transitions for low parameter $q$ values, while predicting abrupt (discontinuous) transitions for high parameter $q$ values.

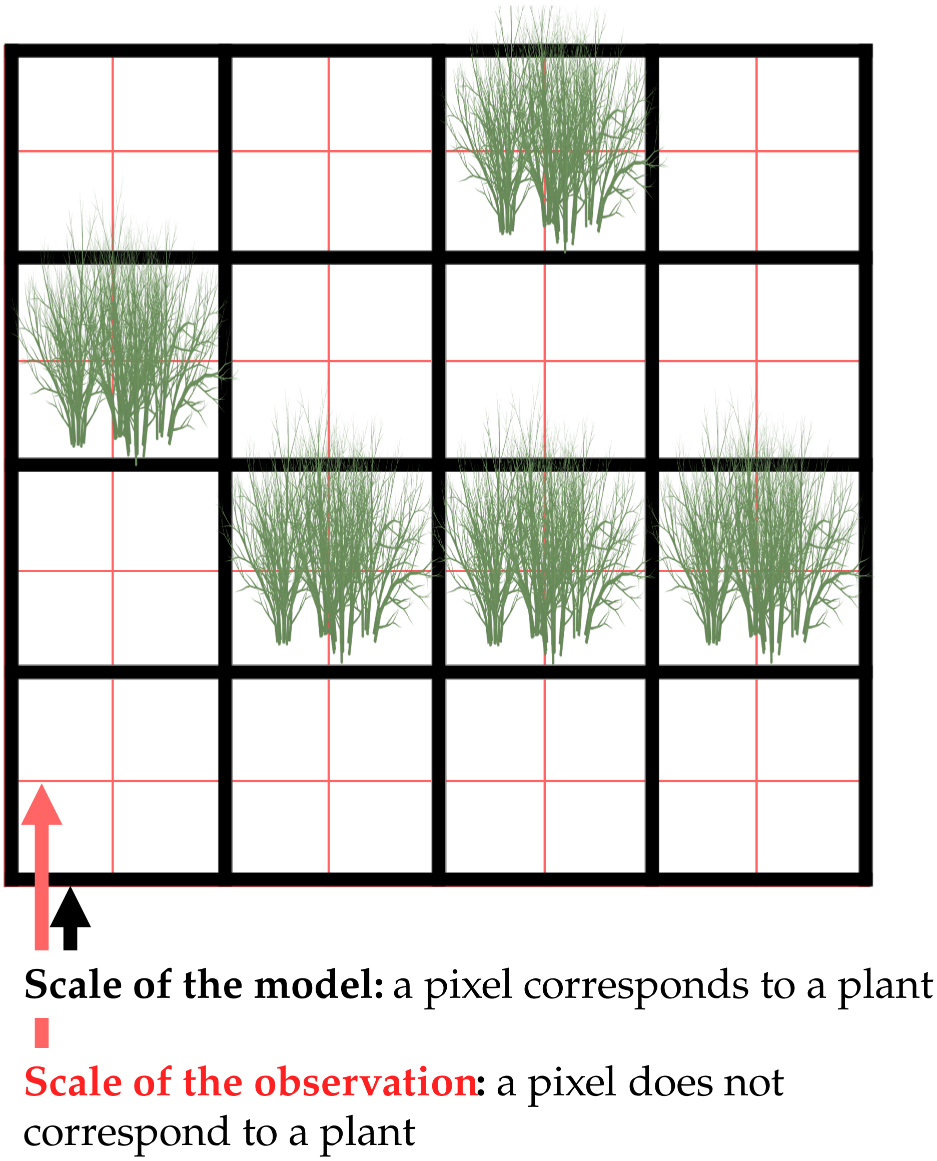


Fig. S15: **Mismatch of scales between simulated and observed vegetation landscapes.** In the model, a pixel is a site colonized or not by a plant. So the pixel is at the spatial scale of plant (*e.g.*, $1m^2$). By contrast, in the observed vegetation landscape, a pixel is a much finer spatial scale (here illustrated for 0.25 $m^2$).

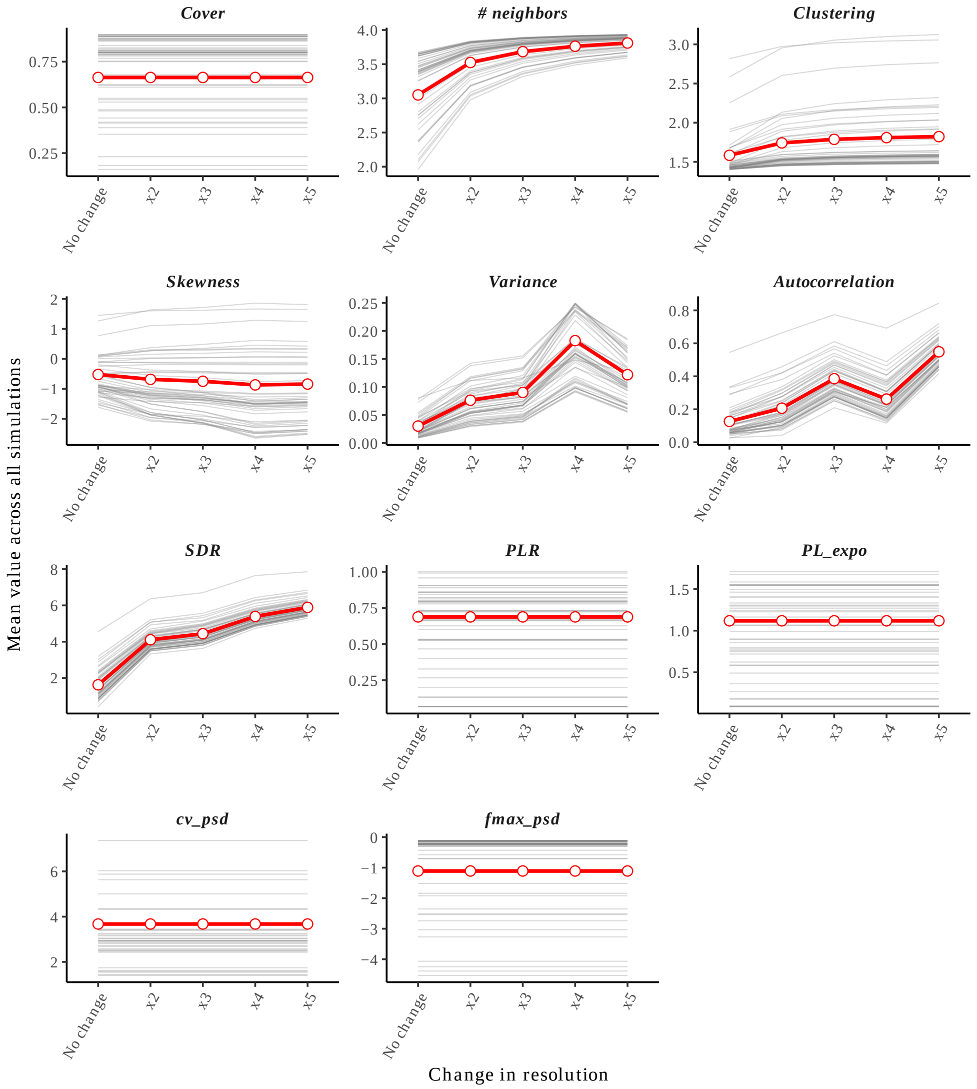


Fig. S16: **Spatial resolution changes the spatial statistics measuring the level of aggregation of the vegetation.**
Using 100 simulated vegetation landscapes, we artificially increased the spatial resolution of the simulations by replacing each pixel with a square of 4 ("x2", $\eta = 2$), 9 ("x2", $\eta = 3$), 16 ("x4", $\eta = 4$) or 25 ("x5", $\eta = 5$) pixels and computed the different spatial statistics. The red line indicates the mean across all simulations, while the gray lines are examples of behavior for 50 landscapes. We observe that despite no change in vegetation cover, accounting for the scale of observation in the model changes the metrics linked with auto-correlation and variance. SDR = Spectral density ratio, Frac. max = Fraction of the landscape covered by the largest patch, CV PSD = coefficient of variation of the patch size distribution.

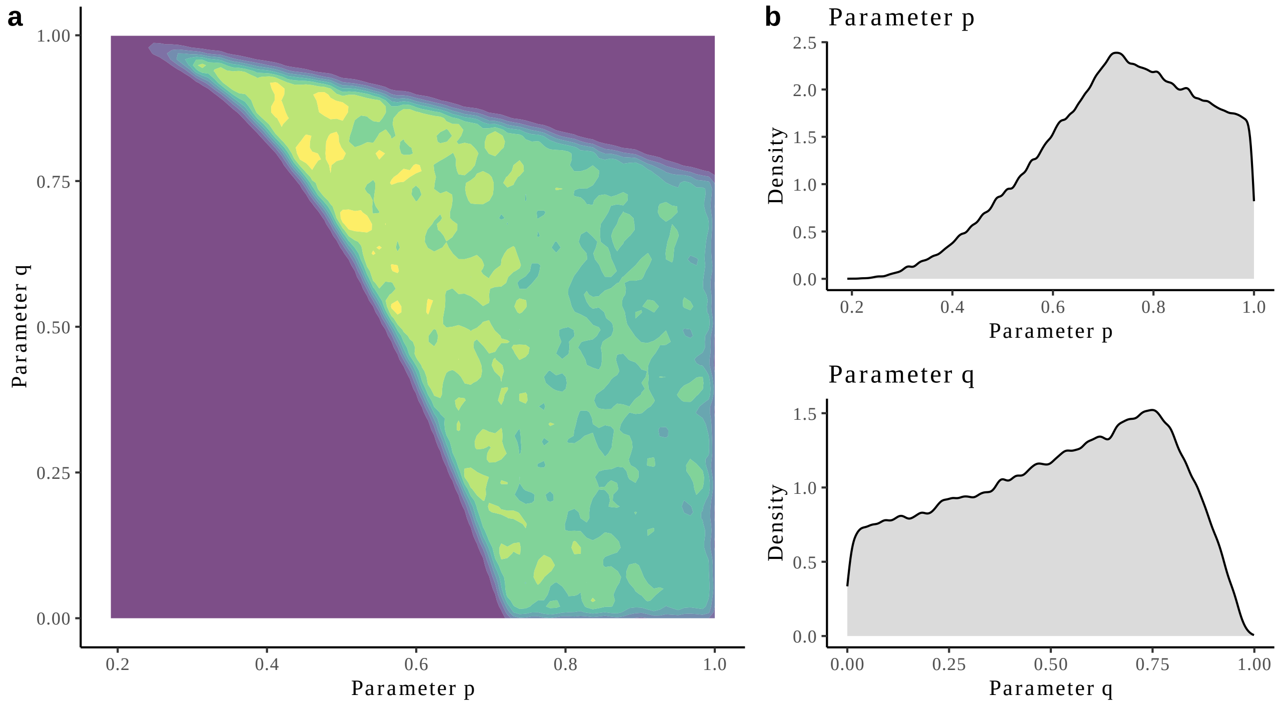


Fig. S17: **Empirical priors of $p$ and $q$.**
Empirical priors of $p$ and $q$. (A) Bi-variate prior density of $p$ and $q$. Color corresponds to the density of simulations: yellow corresponds to high density of sampled parameters, while in purple areas, no parameters were sampled. (B) Density of the prior distribution of $p$ and $q$.

Caption for Fig. S18 **PCA on the 11 spatial statistics reveals model-data gap when not accounting for the scale of observation in the minimal model while accounting for it allows simulations to cover most of the empirical sites.**
We performed a principal component analysis on the 11 spatial statistics computed on both observations and simulations generated by sampling the prior distributions. The 3 first principal components are shown and account for 90.6% of the variance. (Top) We do not account for the mismatch of scales between simulations and observations ($\eta = 1$ by default), which generates a clear gap between the spatial statistics computed on the simulations and the observations. (Bottom) We accounted for the mismatch of scales between simulations and observations by sampling $\eta \in 1, \ldots, 5$ ("Model, x2", $\eta = 2$), 9 ("Model, x3", $\eta = 3$), 16 ("Model, x4", $\eta = 4$) or 25 ("Model, x5", $\eta = 5$), which allows simulations to cover well most of the observations. See legend of Fig. S12 for details on the spatial statistics used.

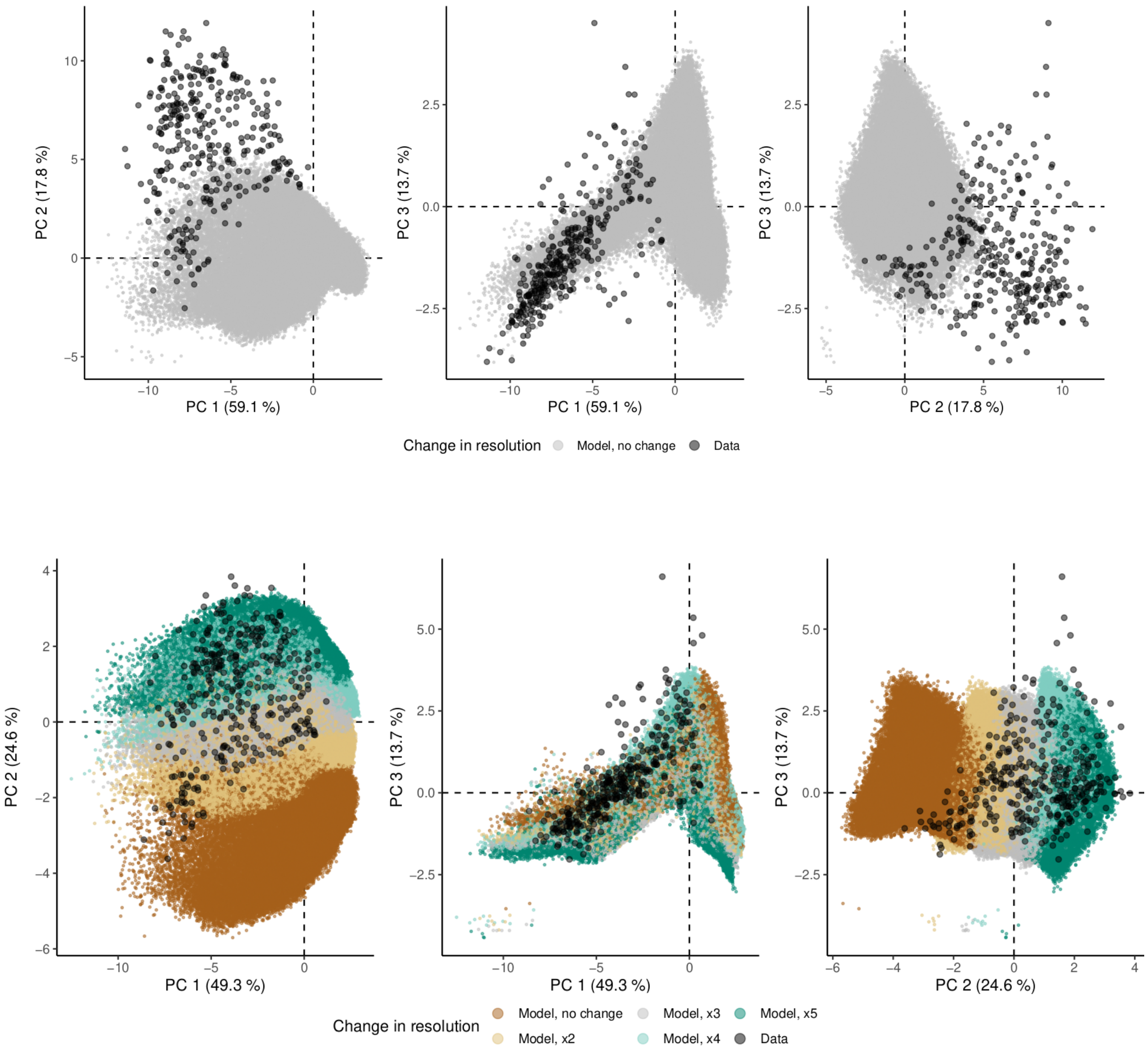


Fig. S18: **Caption is above**

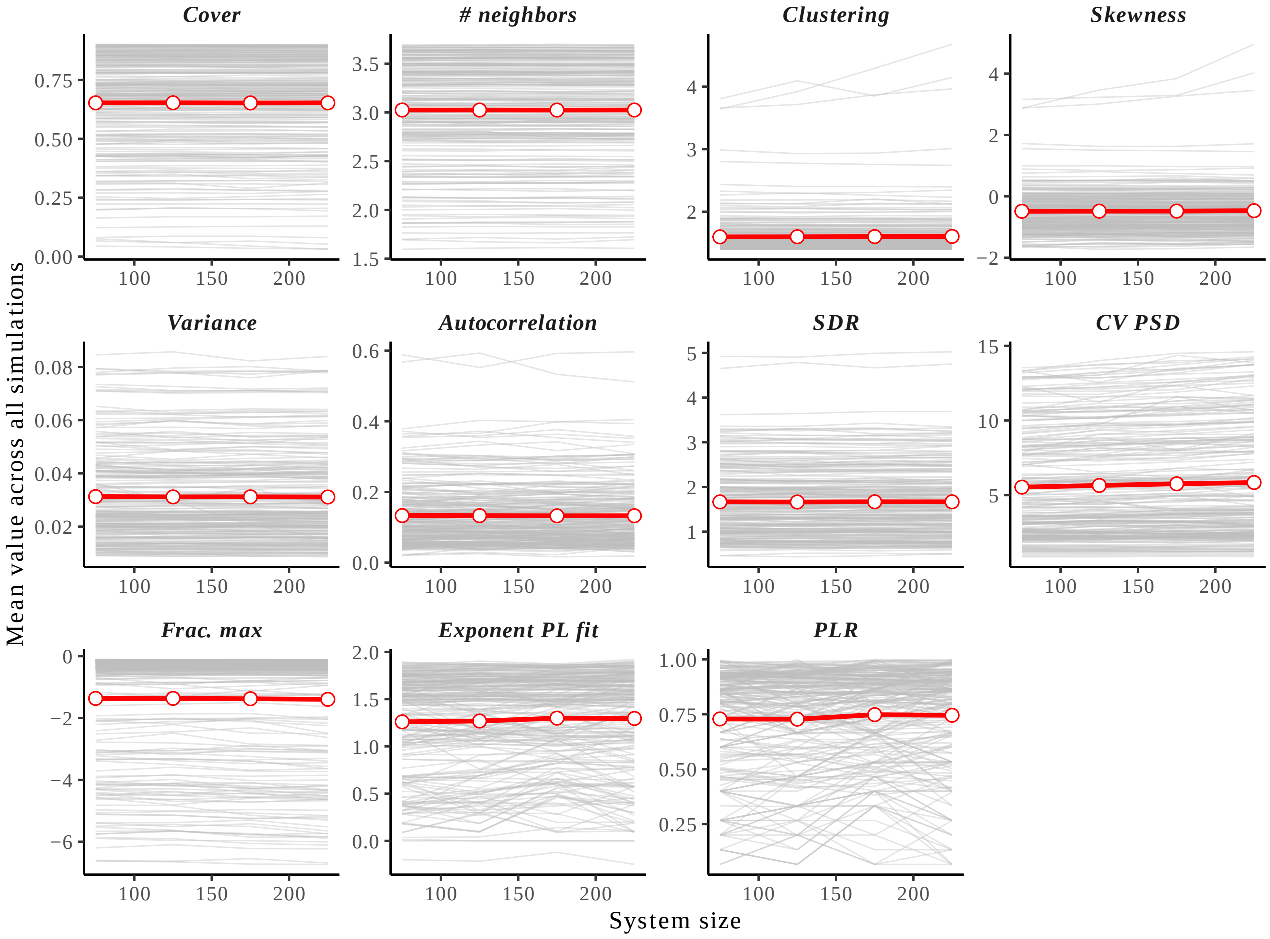


Fig. S19: **Summary statistics do not vary with system size.**
For a subset of 100 randomly chosen parameter values, we ran the minimal model on different landscape sizes (from 75x75 sites to 225x225 sites. Red lines correspond to the average across all parameter values while grey lines correspond to each landscape. This suggests that the spatial statistics we compute do not change with landscape size.

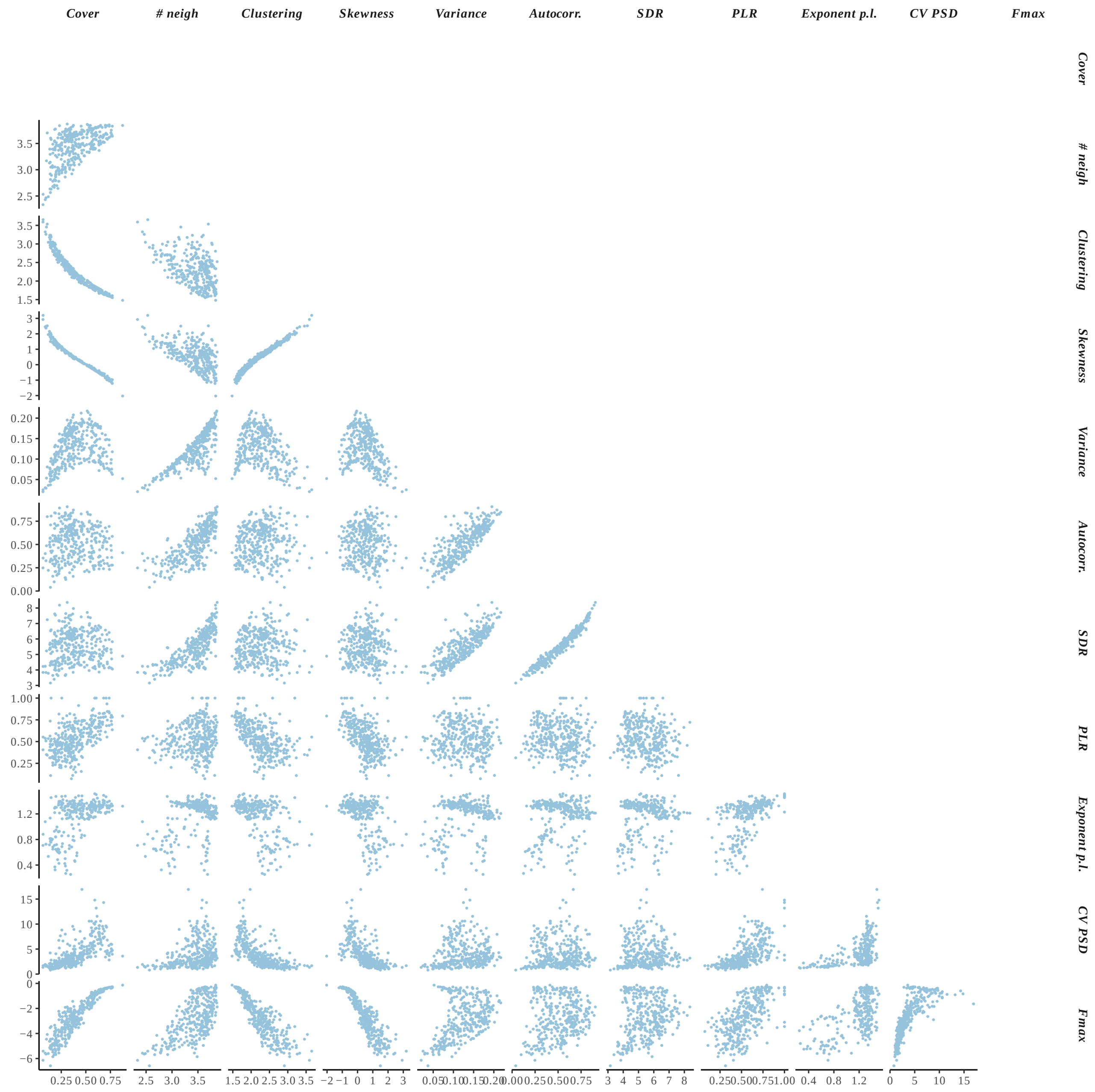


Fig. S20: **Pair correlation between the metrics used to characterize vegetation landscapes.**

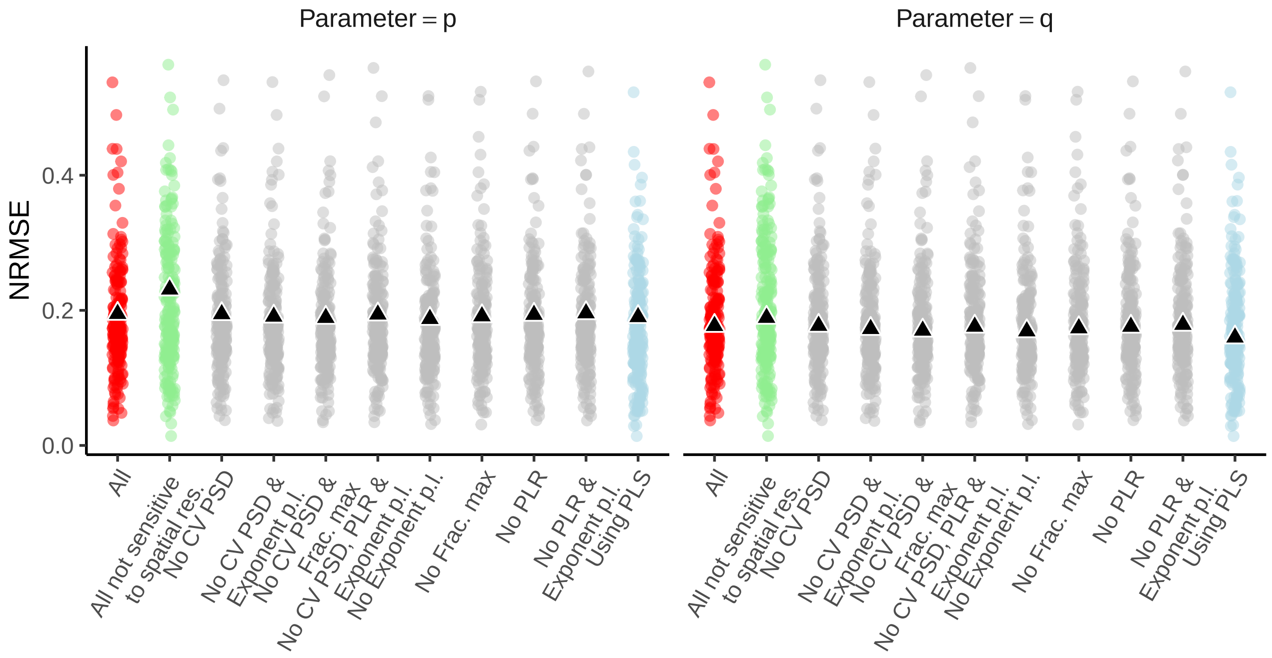


Fig. S21: **Robustness of our inference approach to recover the parameters of simulated landscapes.**
We randomly generated 100 simulated vegetation landscapes (for which the parameters are known). Then, we performed ABC using different subsets of spatial statistics (x-axis) to characterize the spatial structure. We then compared the inferred parameters with the true ones using the NRMSE (see Methods). The red points indicate the case where all statistics have been kept (which we use for main text analysis), while the gray points correspond to inferences with only a subset of all the summary statistics. The blue points indicate the case where we performed partial-least square before the selection of the simulations in order to reduce the dimension of the summary statistics. When performing this prior dimensional reduction, we show that the accuracy on the estimation of parameter values remains similar as for the case where we use all eleven spatial statistics (blue *versus* red points). The green points indicate the case where we only use spatial statistics that are independent of the spatial resolution of the landscape, in which case we still observed a small decrease of the quality of estimation of the parameter values (green *versus* red points).Aside from these two cases, we see no important effects of the subset of spatial statistics used on the ability of the method to recover the parameters of simulated vegetation landscapes. See legend of Fig. S12 for details on the spatial statistics used.

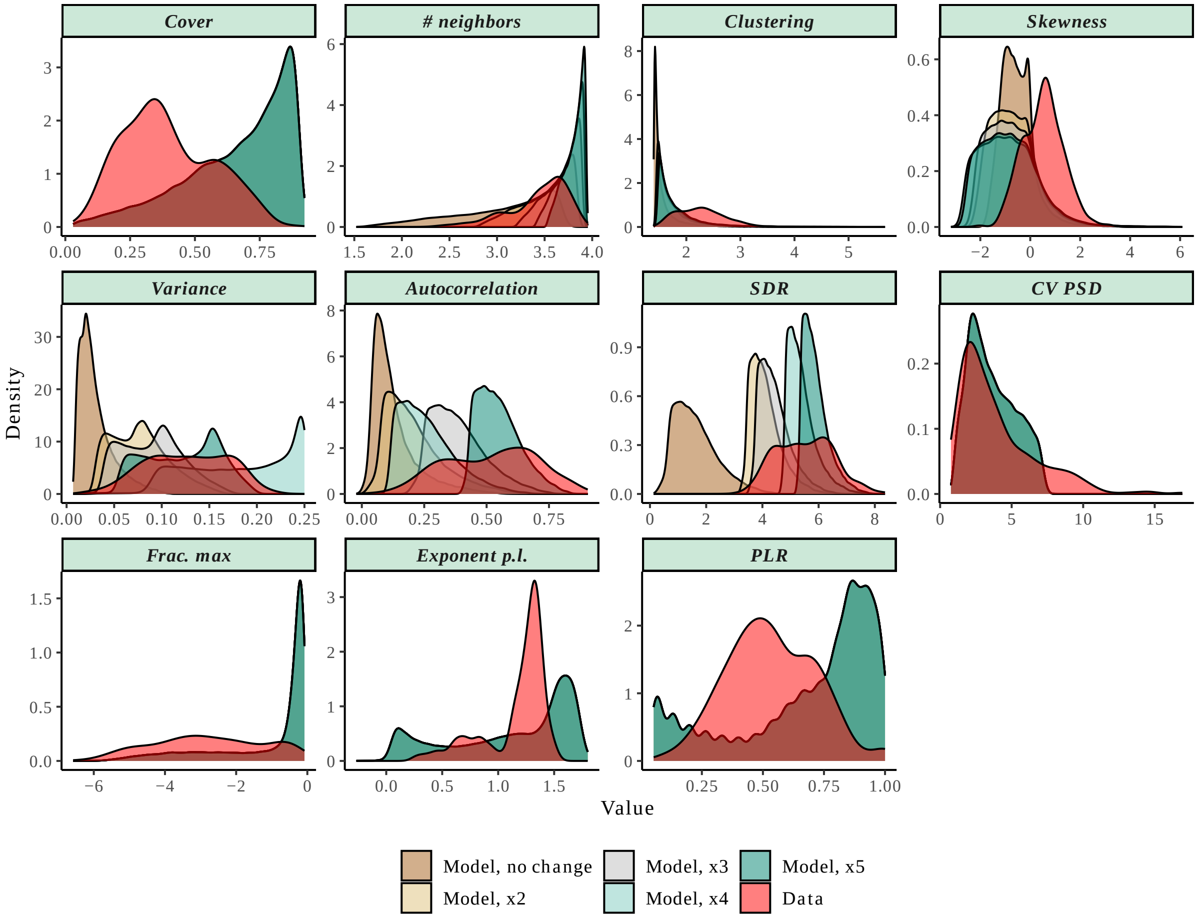


Fig. S22: **The scale layer of the VDmin model allows covering the observations in terms of spatial statistics.**
For each of the 11 spatial statistics, we display the density of the observations as well as the simulations. The color indicated the value of the parameter $\eta$ (no change : $\eta = 1$, $\times 2, \ldots, \times 5 : \eta = 2, \ldots, \eta = 5$). See legend of Fig. S12 for details on the spatial statistics used.

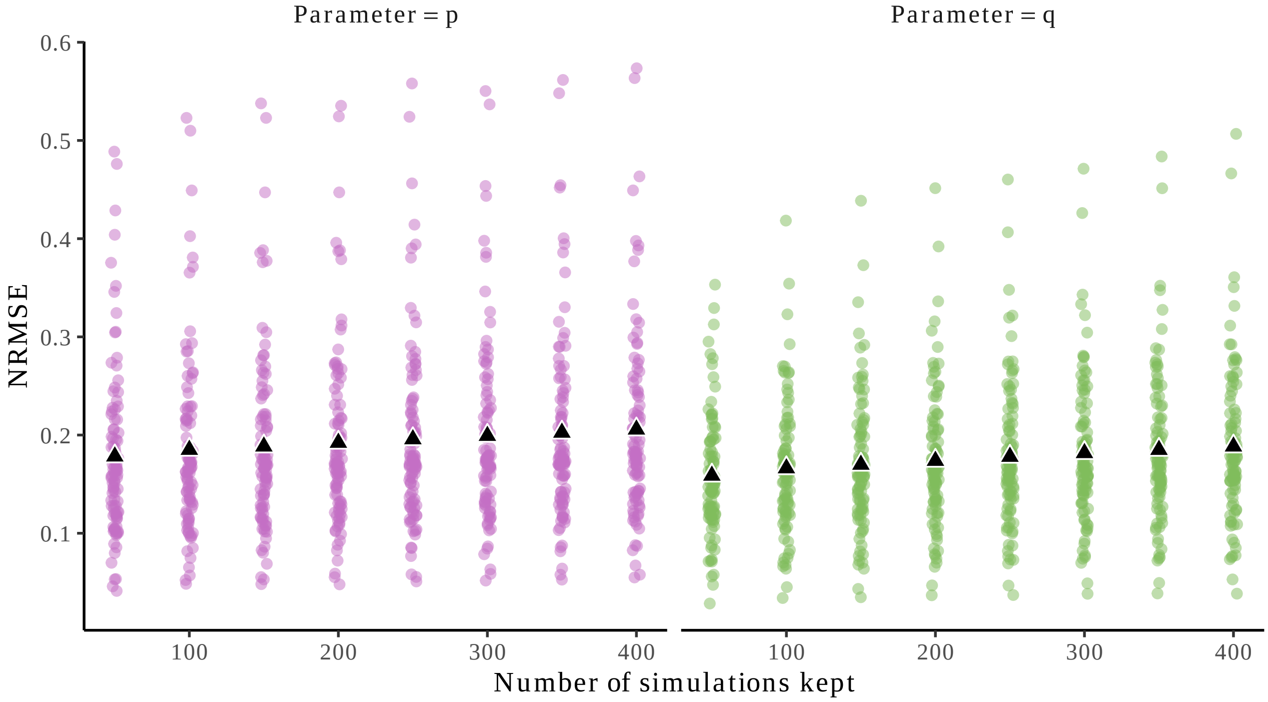


Fig. S23: **Influence of the number of simulations kept on the inference accuracy.** We performed ABC on 100 randomly sampled simulated landscapes (for which the parameters are known) by keeping the 50 . . . 400 closest simulations. We compare the ability of the approach to recover the parameters of these virtual landscapes using the NRMSE $= \frac{\text{RMSE}_{\text{selected}}}{\text{RMSE}_{\text{prior}}}$. As we increased the number of accepted simulations, bias on the posterior distribution of $p_{est}$ and $q_{est}$ increased. We kept 100 accepted simulations *(i)* following (42) who accepted 0.0001% of the simulations and *(ii)* as a trade-off between the inference accuracy and the number of simulations kept to build the posterior distribution $\theta^*$.

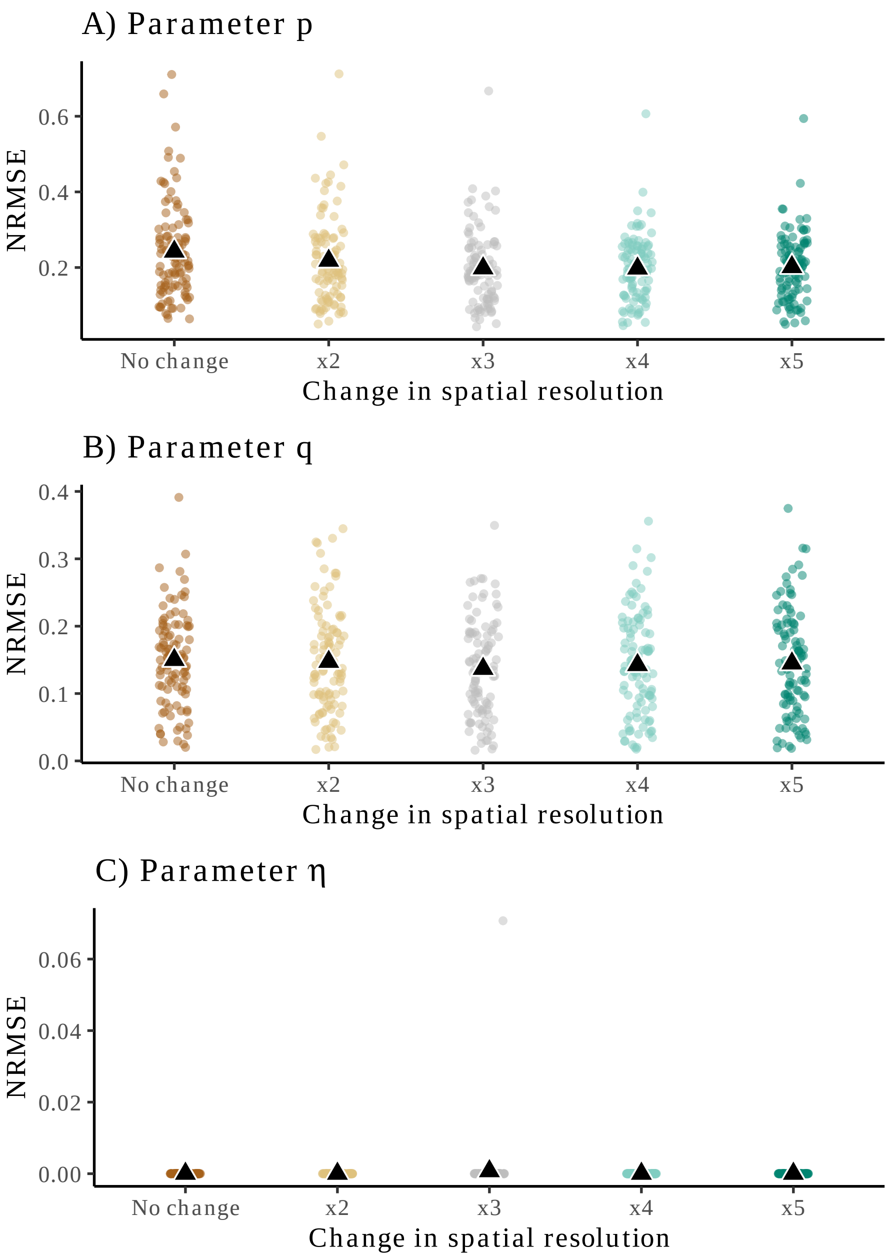


Fig. S24: **The scale of observation is well recovered when performing ABC.** We compare the ability of the ABC approach to recover the true parameters of $N = 100$ simulations (virtual landscapes) and for which we varied $\eta$ (*i.e.,* different scale parameter). To evaluate the quality of the inference, we computed the NRMSE (see Methods). When NRMSE is close to 0, we retrieve very precisely the parameter of the virtual landscapes. For $p$ (A; mean NRMSE = 0.211), $q$ (B, mean NRMSE = 0.144), and $\eta$ (C, mean NRMSE = 0.00014) the NRMSE is very low meaning that we accurately infer the parameters of simulated landscapes.

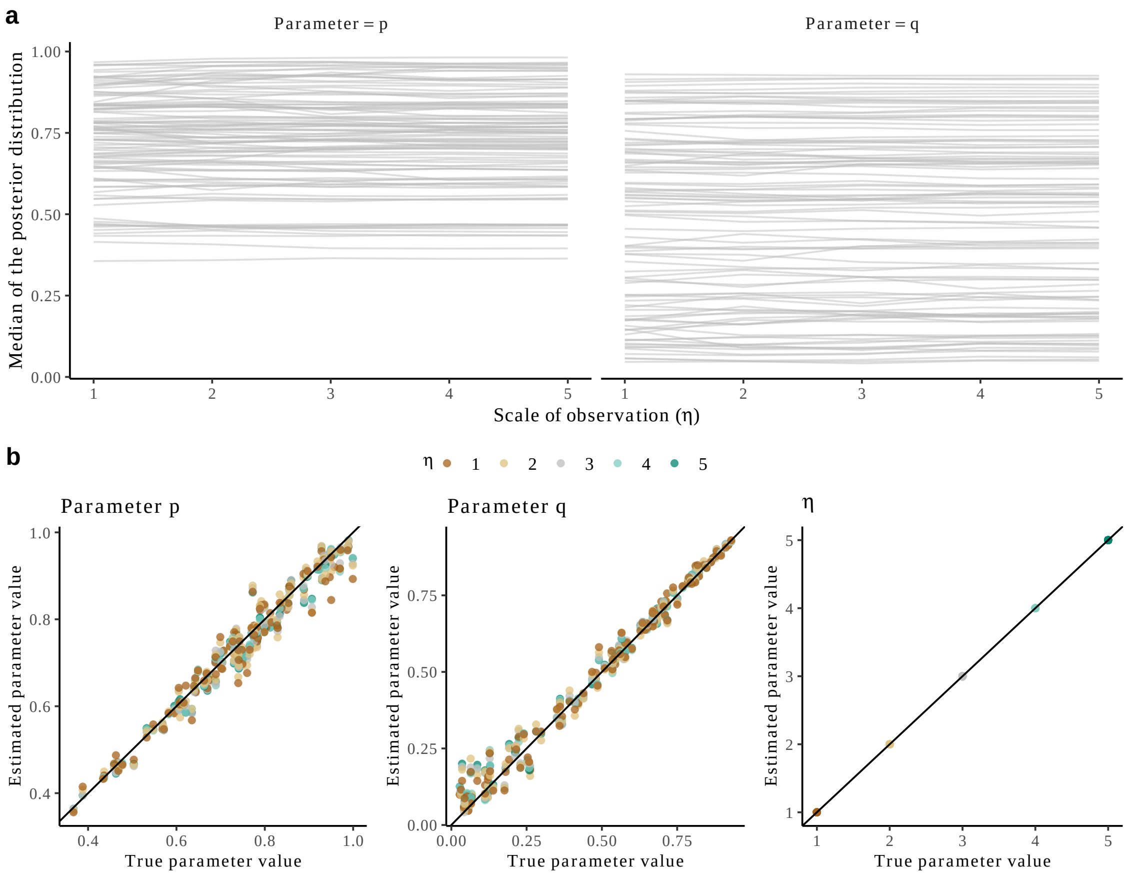


Fig. S25: **The scale layer does not change the parameters inferred by the approach.** We compare the ability of the approach to recover the parameters $(p, q)$ of $N = 100$ virtual landscapes observed at different scales (*i.e.,* characterized by different spatial resolution, x-axis). (A) We see that no matter the parameter of the scale layer, we recover the same mean posterior of $p$ and $q$ for each of the 100 landscapes even if each can be characterized by different spatial resolutions' parameter $\eta$. Note that observed deviations from a strict horizontal lines are due to model stochasticity and non-infinite sampling of the parameter space. (B) Same data but represented by comparing true parameter values used to run the simulations and estimated ones using the ABC. The color indicates the parameter $\eta$ and shows that the bias is similar for the different values of $\eta$.

Therefore, no matter the spatial resolution in the data, accounting for the spatial resolution in the model allows us to recover the parameters associated with each landscape structure.

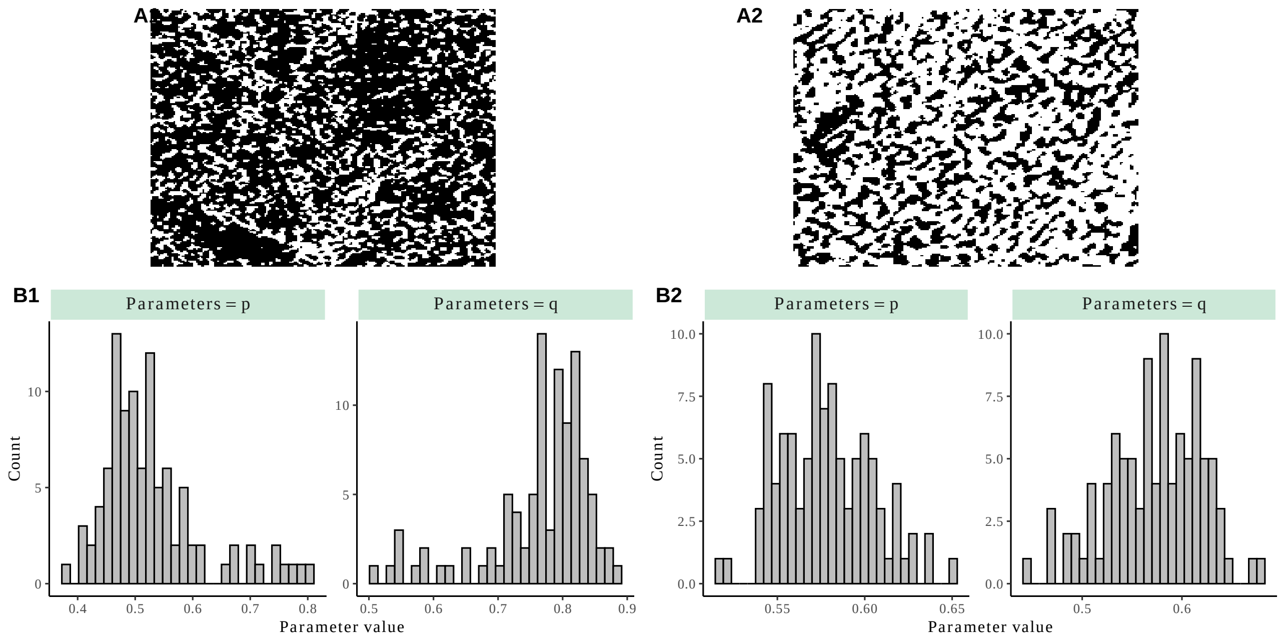


Fig. S26: **Examples of the posterior distributions of the two parameters ($p_{est}$,$q_{est}$) for two sites with contrasted spatial structure.**
These are sites 170-b and 116-c from the BIOCOM dyland dataset (2).

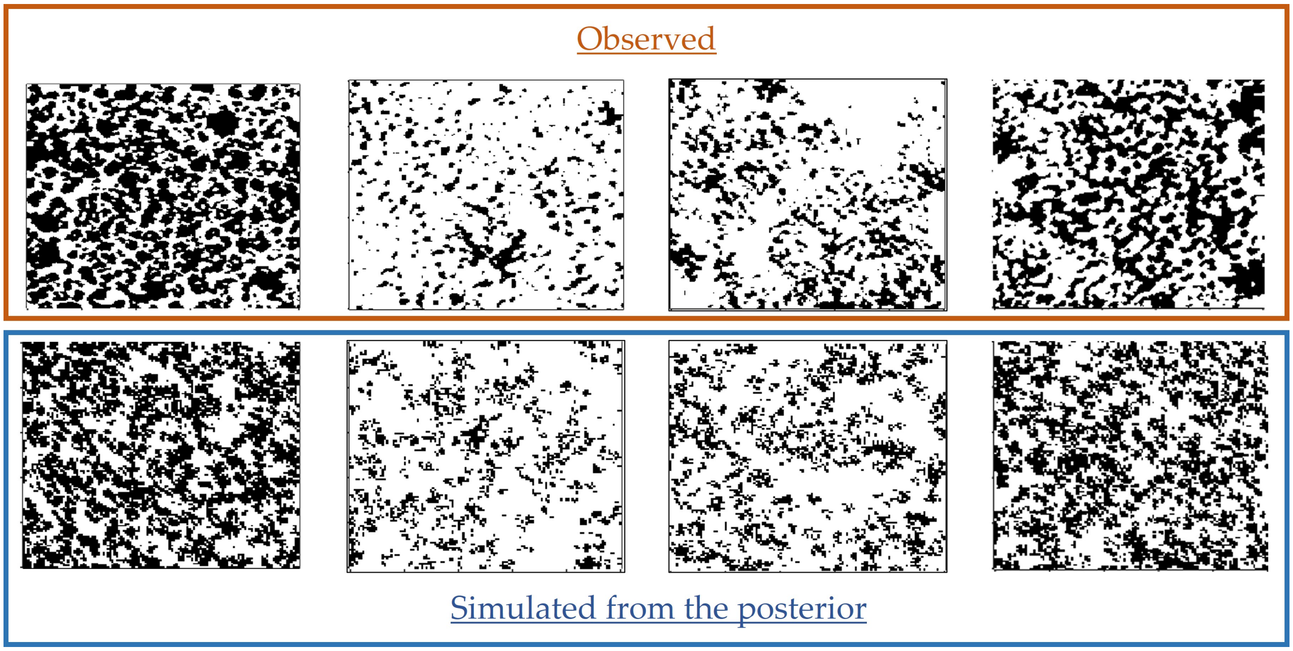


Fig. S27: **Illustration that simulated landscapes generate similar spatial structures compared to the observed ones.**
For each of the four observed vegetation landscapes, we sampled a couple of parameters $(p, q)$ from the posterior distribution of the parameters and ran the model (*i.e.*, posterior predictive checks). A landscape with at the asymptotic state is shown (bottom row), as well as its corresponding observation (top row).

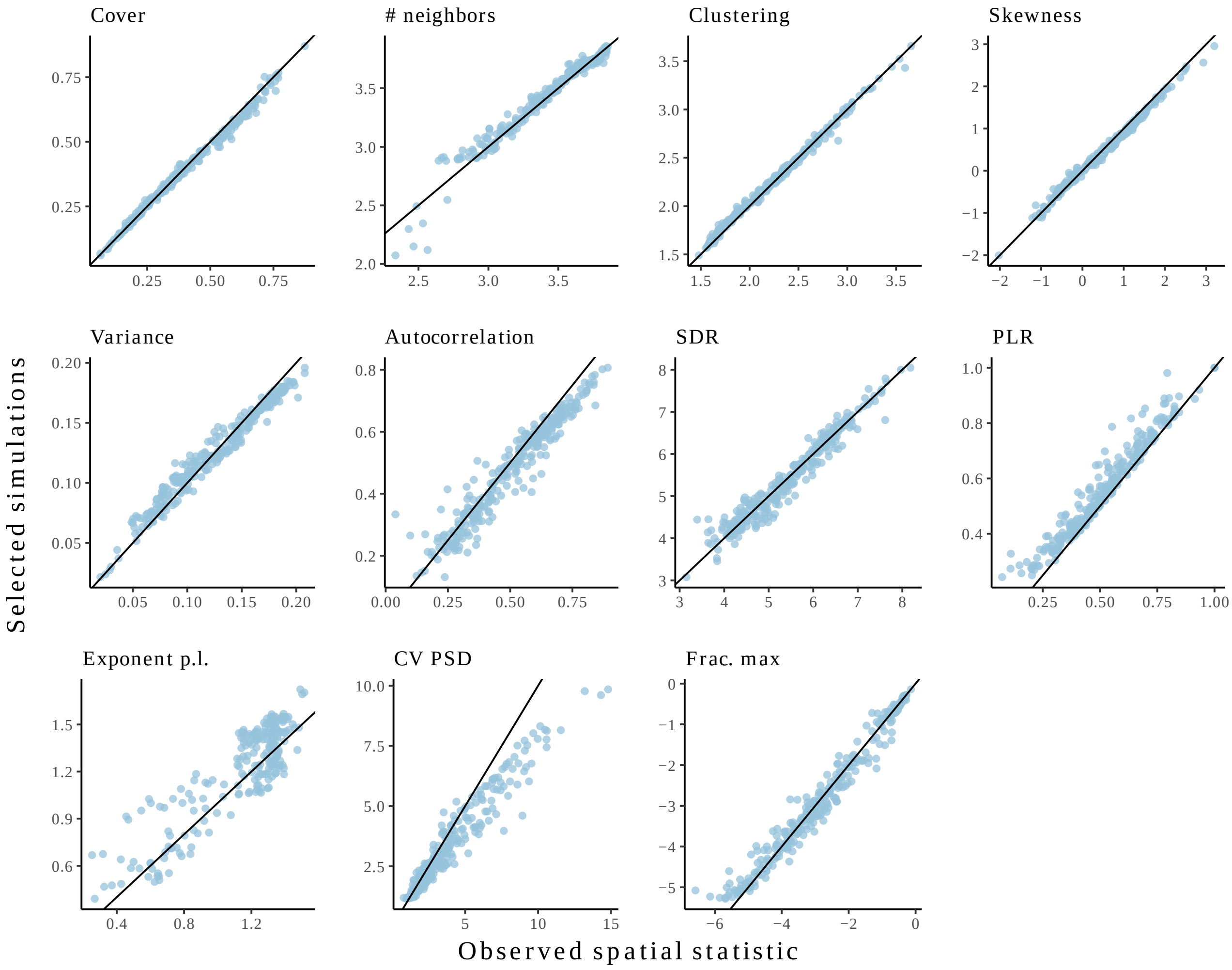


Fig. S28: **Simulations recover well the level of aggregation, vegetation cover and fits on the patch-size distribution but lack of heterogeneity in patch size compared to observations.**
We performed ABC on each of the 293 observed landscapes and here compare the mean spatial statistics of the 250 closest simulations for each observation (y-axis) with the observed spatial statistic (x-axis). While simulations perform well at replicating the level of vegetation aggregation in the data (variance, clustering, auto-correlation, or SDR metrics), they perform less well when there is a high heterogeneity in the patch size distribution of observed landscapes is high (*i.e.,* high CV PSD). As an example, see the comparison between observed and simulated landscape in Fig. S26. See legend of Fig. S12 for details on the spatial statistics used.

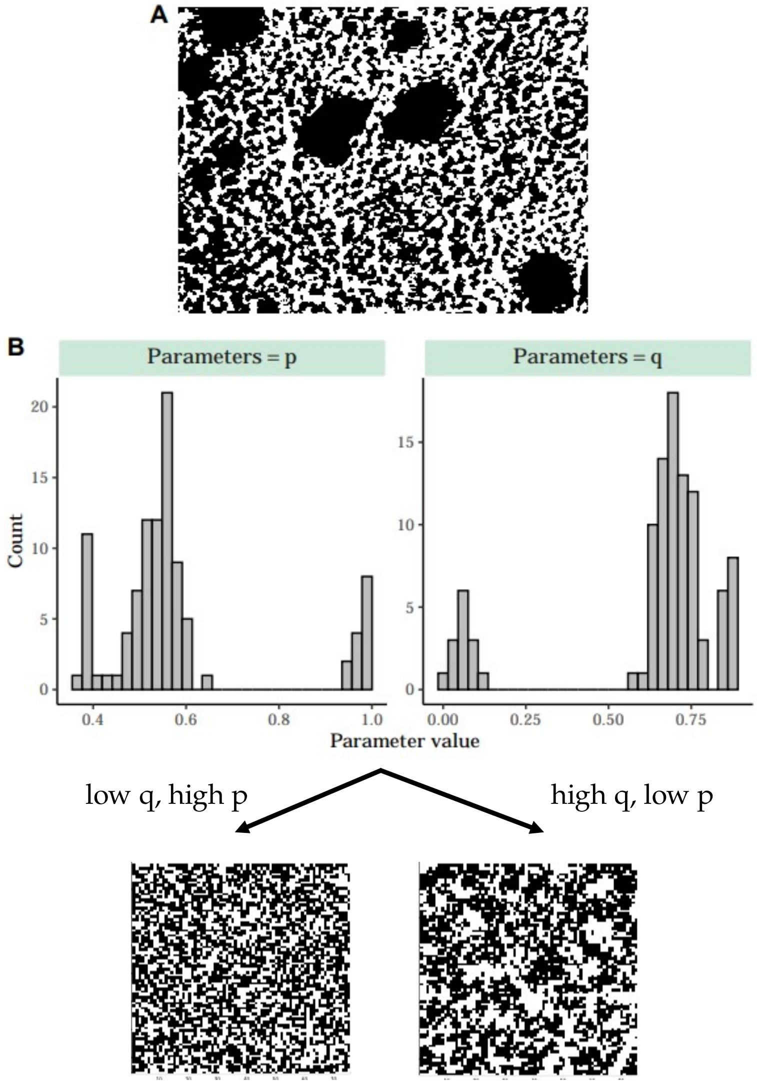


Fig. S29: **Example of a site where contrasted scales lead to bimodal posterior distributions of $p$ and $q$.**
(A) Example of a binary vegetation landscape removed from the analysis. There are contrasting scales with coexistence of small grass species (likely) and trees. (B) Consequently, in the posterior distribution of $p_{est}$ and $q_{est}$, there is a clear bimodality that emerges due to the strong heterogeneity in the patch size distribution. Consequently, because the model lacks heterogeneity compared to the data, the ABC method selects simulations that generate either large patches (upper tail of the distribution of $q_{est}$, right landscape on the bottom) or smaller patches (lower tail of the distribution of $q_{est}$, left landscape on the bottom).

Caption for Fig. S30. **When the spatial resolution gets too low, the observed spatial structure is blured, the spatial statistics from the closest selected simulations gets more and more different to the observed ones, and the inferred parameters change.**

(A) Comparison between the spatial statistics of the closest accepted simulations and the ones computed on the observed ecosystem images. To evaluate how closely the spatial structure in the simulations and the observations (here virtual observations) are, we computed the NRMSE (see Methods) for n=100 virtual observations (corresponding to the points). When the NRMSE increases to 1 (black line), the average spatial statistics in the closest selected simulations are more and more different compared to the observed ones. We compare the ecosystem images with their initial spatial resolution ("No change"), with a spatial resolution two-times lower ("Resolution /2"), and with a spatial resolution three-times lower ("Resolution /3") using coarse-graining. The colored points correspond to each of the 293 ecosystem images and the black point is the mean.

(B) Change of the median of the posterior distribution of parameter $p$, $q$ and $\eta$ for the different spatial resolution of the ecosystem images. Each line grey corresponds to one of the 293 ecosystem images, while the red line corresponds to the mean.

When spatial resolution is low (*e.g.,* for "Resolution /3"), the size the pixels can be larger than a single plant individual (as explained in Methods), which therefore breaks the fine-scale spatial structure of the vegetation, leads to a poorer fit with the simulations (panel A) and to important changes in the posterior distribution of the parameter $p$ and $q$ (panel B).

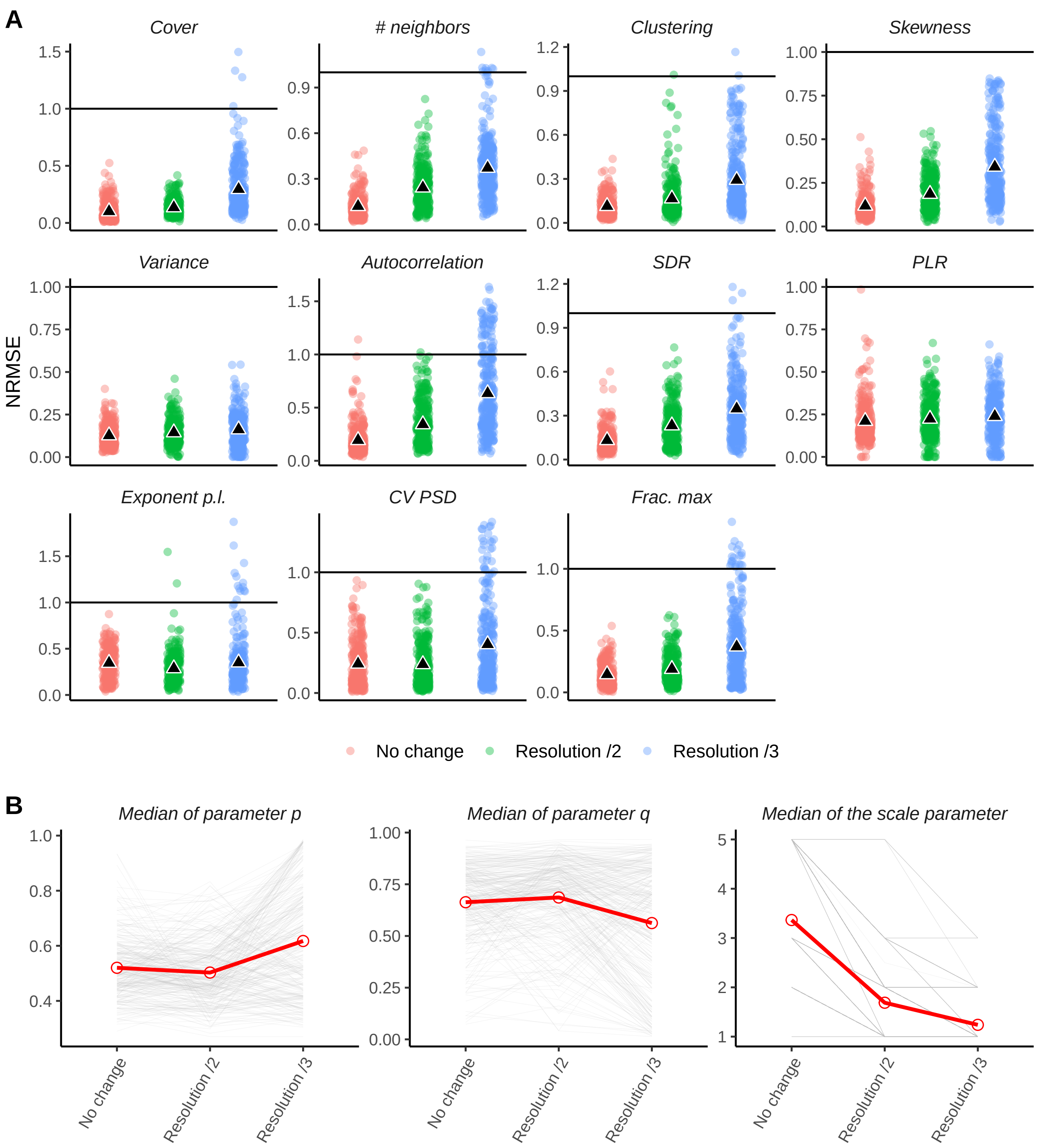
A
Cover
### neighbors
Clustering
Skewness
Variance
Autocorrelation
SDR
PLR
Exponent p.l.
CV PSD
Frac. max
NRMSE
No change
Resolution /2
Resolution /3
B
Median of parameter p
Median of parameter q
Median of the scale parameter

Fig. S30:

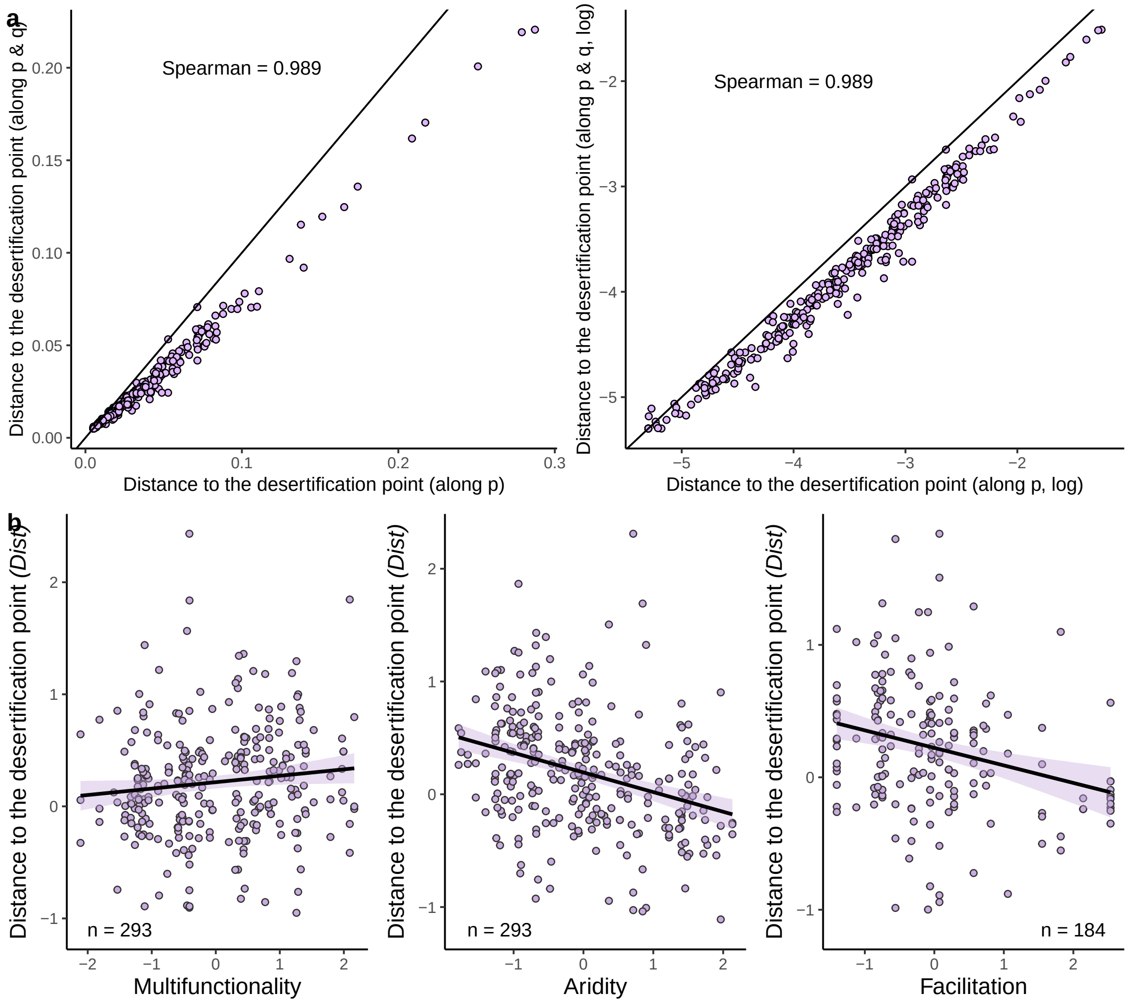


Fig. S31: **Distance to the desertification point estimated by decreasing $p$ and $q$ leads to similar results compared to the distance estimated by decreasing $p$.**
(a) Comparison between the median of the posterior distribution of the estimated distance to the desertification point when varying $p$ alone (x-axis) and decreasing both $p$ and $q$ simultaneously (y-axis). The distances are log-transformed in the right pannel compared to the left one. (b) Same analysis as in Fig.3a but using the distance estimated by decreasing both $p$ and $q$ simultaneously. The points show partial residuals, and the lines correspond to model fits. Shading around each line represents the 95% confidence interval.

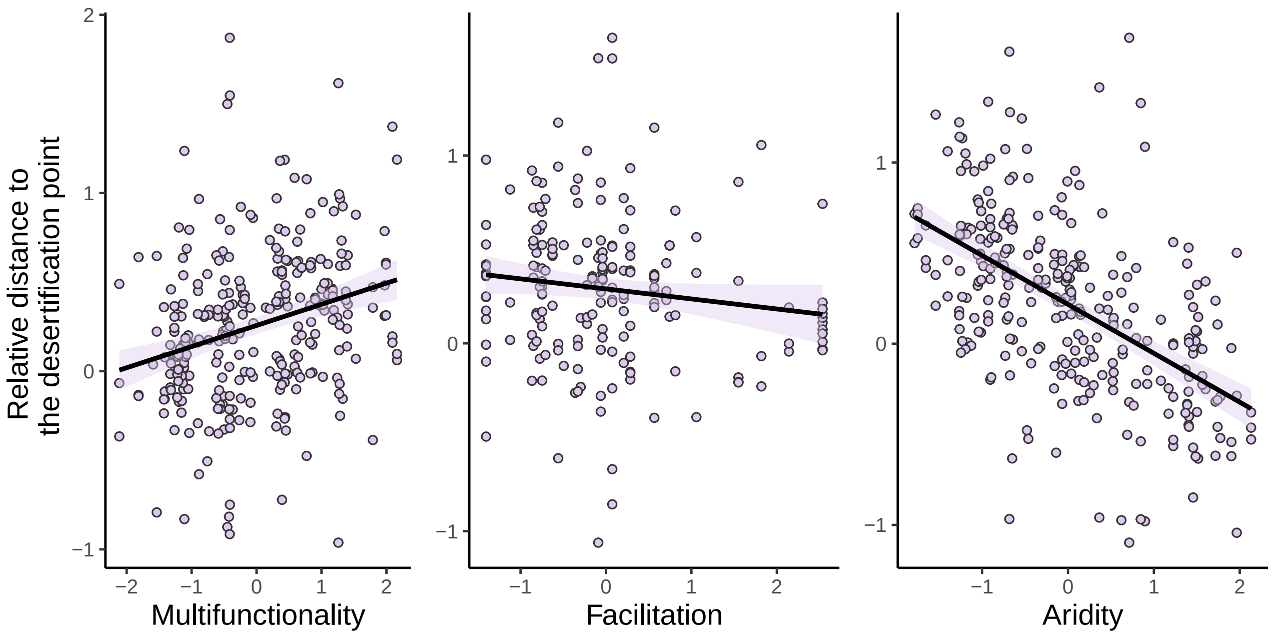


Fig. S32: **Drivers of the estimated distance to the desertification point of global drylands using the relative distance to the desertification point.**
(A) Predicted responses of the relative distance of the desertification point to changes in aridity, multifunctionality and level of facilitation. The points show partial residuals, and the lines correspond to model fits. Shading around each line represents the 95% confidence interval.

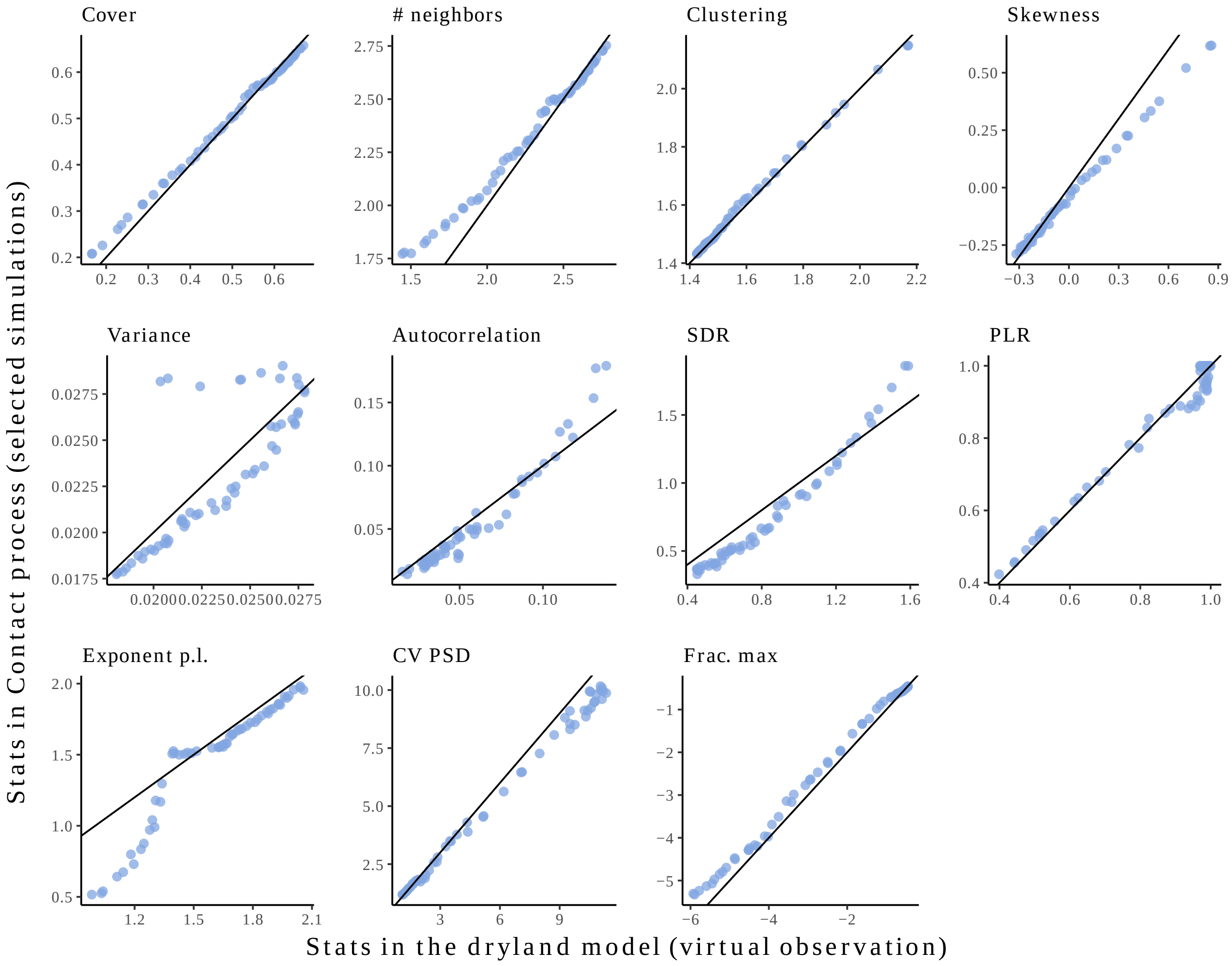


Fig. S33: **The VDmin model covered well the spatial statistics of the landscape of vegetation from the Kéfi model.**

We performed our inference approach on each of the virtual vegetation landscapes simulated from the different parameter sets of the Kéfi model (*16*). We compare the mean spatial statistics of the 100 closest simulations (y-axis), with the observed spatial statistic (x-axis). See for the parameter values used for the Kéfi model (”Validating our approach using other models”).

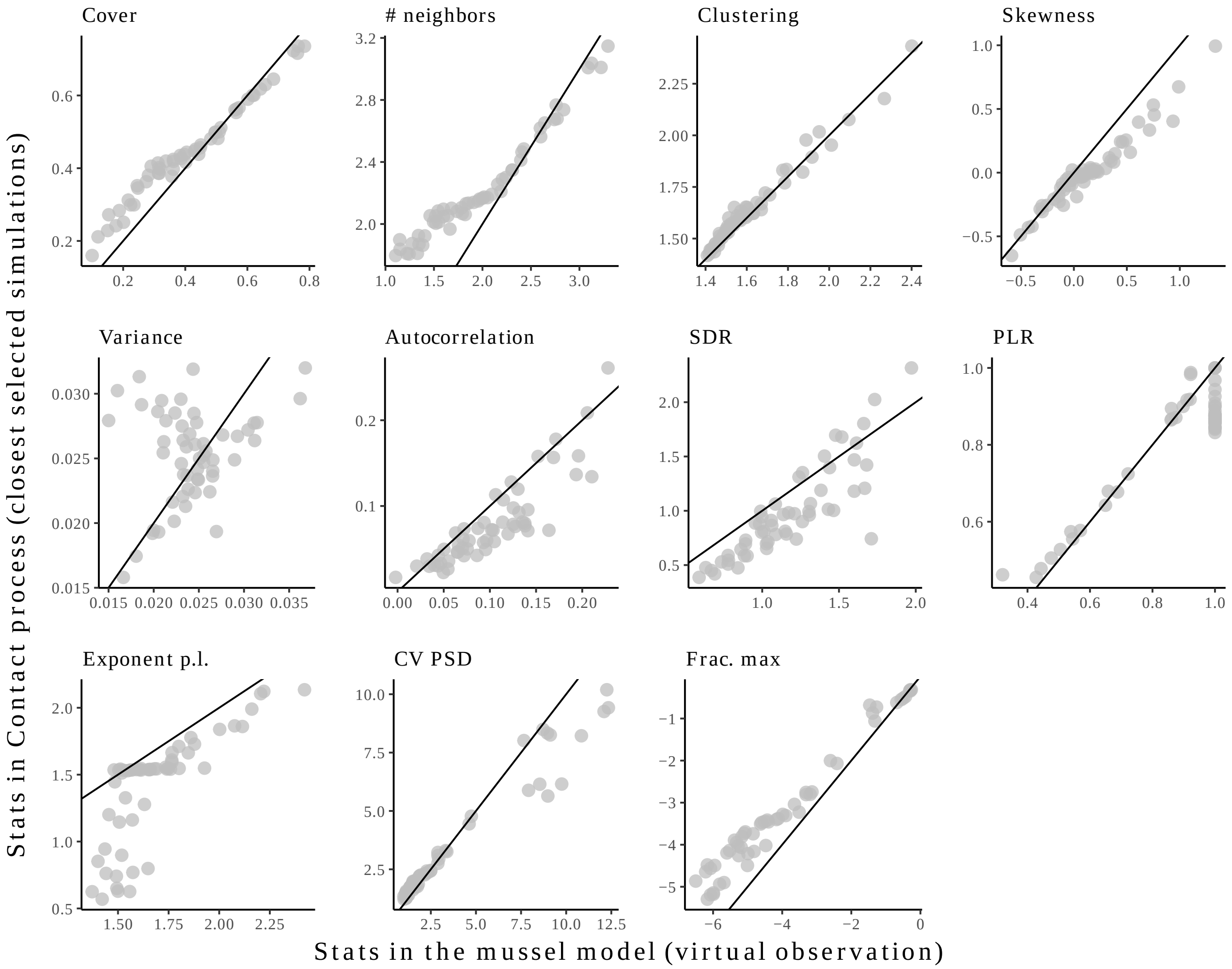


Fig. S34: **The VDmin model covered well the spatial statistics of the landscape of vegetation from the Guichard model**
We performed our inference approach on each of the virtual vegetation landscapes simulated from the different parameter sets of the Guichard model (*24*). We compare the mean spatial statistics of the 100 closest simulations (y-axis), with the observed spatial statistic (x-axis). See legend of Fig. S12 for details on the spatial statistics used. See Methods for the definition of the spatial statistics used ("Choice of the discrepancy between observations and simulations"), and for the parameter values used for the Guichard model ("Validating our approach using other models").

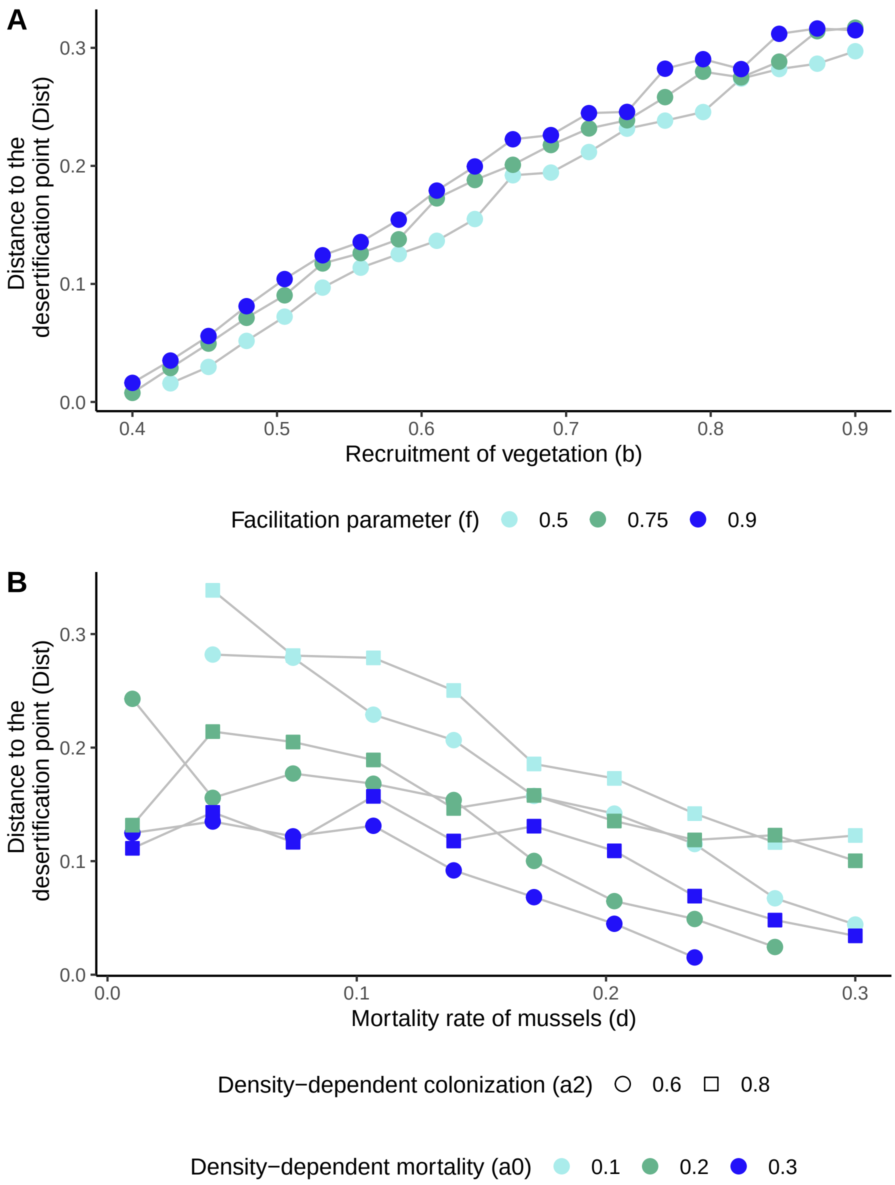


Fig. S35: **Linking parameters in the two original models with the estimated distance to the degradation point.**
Changes in estimated distance to the degradation point along model parameters used for (a) the dryland vegetation model and (b) the mussel-bed models.

### Supplementary tables (S1-S3)

Table S1: **Results of the statistical model presented in Eq. S14 for the estimated distance to the desertification point (*Dist*) (Fig. 3A-C).** The marginal and conditional $R^2$, the predictor estimates, their standard errors are indicated. We also indicate the variance inflation factors (VIF) for each predictor, and the results of Moran tests for spatial autocorrelation (performed at different spatial scales, with the 10, 30, and 50 closest plots). The VIF values obtained (typically $< 10$) in all cases suggest that multicollinearity was weak in the model. In addition, there was no spatial autocorrelation in the model.

| Model for the distance to the desertification point. $R^2_m$ = 0.190, $R^2_c$ = 0.718 |
|---|
| Moran tests: k=10 (p=0.994), k=30 (0.893), k=50 (0.807) |

| **Predictor** | **Effect size** | **Std error** | **Vif** |
|---|---|---|---|
| *Aridity* | -0.21 | 0.10 | 1.61 |
| *Multifunctionality* | 0.07 | 0.10 | 1.85 |
| *Sand* | -0.09 | 0.11 | 2.12 |
| *Latitude* | 0.16 | 0.10 | 1.44 |
| *Longitude (cos)* | -0.12 | 0.08 | 1.07 |
| *Longitude (sin)* | 0.01 | 0.08 | 1.15 |
| *Slope* | -0.05 | 0.09 | 1.36 |
| *Elevation* | -0.19 | 0.09 | 1.36 |

Table S2: $R^2$ **(top) and bootstrapped p-values (bottom) of the linear predictions using partial residual plots of Figs. 3.**

| Predictor | Absolute distance | Relative distance | Parameter q | Cover |
|---|---|---|---|---|
| Aridity | 0.169 | 0.289 | 0.103 | 0.621 |
| Multifunctionality | 0.024 | 0.08 | 0.091 | 0.373 |
| Facilitation by nurses | 0.058 | 0.018 | 0.282 | 0.217 |

| Predictor | Absolute distance | Relative distance | Parameter q | Cover |
|---|---|---|---|---|
| Aridity | <0.001 | <0.001 | <0.001 | <0.001 |
| Multifunctionality | <0.001 | 0.003 | <0.001 | <0.001 |
| Facilitation by nurses | <0.001 | 0.012 | <0.001 | <0.001 |

Table S3: **Bootstrapped coefficients and p-values of direct and total paths of the SEM in Fig. 3D.** CI = confidence interval at 95%, Moran I = Spatial autocorrelation of the vegetation.

| Predictor | Response | CI | p-value | Type path |
|---|---|---|---|---|
| *Cover* | Distance | 0.47(0.58,0.35) | $< 0.01$ | Direct |
| *Moran I* | Distance | -0.44(-0.55,-0.32) | $< 0.01$ | Direct |
| *Multifunctionality* | Distance | 0.07(-0.01,0.16) | 0.05 | Total |
| *Aridity* | Distance | -0.21(-0.28,-0.15) | $< 0.01$ | Total |
| *Aridity* | Moran I | 0.17(0.02,0.29) | $< 0.01$ | Direct |
| *Multifunctionality* | Moran I | 0.15(0.02,0.29) | $< 0.01$ | Direct |
| *Aridity* | Cover | -0.35(-0.45,-0.25) | $< 0.01$ | Direct |
| *Multifunctionality* | Cover | 0.30(0.18,0.42) | $< 0.01$ | Direct |
| *Aridity* | Multifunctionality | -0.43(-0.59,-0.21) | $< 0.01$ | Direct |
